# Leaky ribosomal scanning enables tunable translation of bicistronic ORFs in green algae

**DOI:** 10.1101/2024.07.24.605010

**Authors:** Marco A. Duenas, Rory J. Craig, Sean D. Gallaher, Jeffrey L. Moseley, Sabeeha S. Merchant

## Abstract

Advances in sequencing technology have unveiled examples of nucleus-encoded polycistronic genes, once considered rare. Exclusively polycistronic transcripts are prevalent in green algae, although the mechanism by which multiple polypeptides are translated from a single transcript is unknown. Here, we used bioinformatic and in vivo mutational analyses to evaluate competing mechanistic models for polycistronic expression in green algae. High-confidence manually curated datasets of bicistronic loci from two divergent green algae, *Chlamydomonas reinhardtii* and *Auxenochlorella protothecoides*, revealed 1) a preference for weak Kozak-like sequences for ORF 1 and 2) an underrepresentation of potential initiation codons before ORF 2, which are suitable conditions for leaky scanning to allow ORF 2 translation. We used mutational analysis in *Auxenochlorella protothecoides* to test the mechanism. In vivo manipulation of the ORF 1 Kozak-like sequence and start codon altered reporter expression at ORF 2, with a weaker Kozak-like sequence enhancing expression and a stronger one diminishing it. A synthetic bicistronic dual reporter demonstrated inversely adjustable activity of green fluorescent protein expressed from ORF 1 and luciferase from ORF 2, depending on the strength of the ORF 1 Kozak-like sequence. Our findings demonstrate that translation of multiple ORFs in green algal bicistronic transcripts is consistent with episodic leaky ribosome scanning of ORF 1 to allow translation at ORF 2. This work has implications for the potential functionality of upstream open reading frames found across eukaryotic genomes and for transgene expression in synthetic biology applications.

**Significance Statement:** Textbook dogma states that nucleus-encoded genes are monocistronic, producing transcripts with a single translated open reading frame. However, highly conserved bicistronic loci are pervasive in green algae that are separated by several hundred million years of evolution, speaking to their ancestral origins and functions within the Chlorophyte lineage. A combination of bioinformatic analysis and in vivo gene manipulation supports leaky ribosomal scanning as the primary mechanism for translation of multiple ORFs from bicistronic transcripts. We have successfully tuned synthesis levels of two proteins encoded on one mRNA by modifying the ORF 1 Kozak-like sequence. These findings may have broad applications in synthetic biology.

## Introduction

Eukaryotic translation initiation begins at the 5’ cap, where eukaryotic initiation factors (eIFs), the initiator tRNA, and the 40S ribosomal subunit converge to form the 43S pre-initiation complex (1). Once assembled, the pre-initiation complex scans downstream, seeking a suitable translation initiation site. This is generally discerned by the presence of an initiator codon (AUG) and its surrounding sequence context (2). The optimum sequence, referred to as the Kozak consensus, impacts the efficiency of recognition of the initiator codon by the pre-initiation complex, and features motifs conserved throughout eukaryotes (3–5). After identifying an appropriate initiator codon, 80S assembly, polypeptide elongation, termination, and ribosome disassembly proceed (6). The cap-dependent nature and linear directional scanning of the pre-initiation complex typically prioritizes the first start codon encountered, limiting the recognition of potential downstream ORFs (7, 8). Consequently, eukaryotic nuclear genes have traditionally been considered monocistronic; encoding mature mRNAs that contain a single protein-coding ORF.

Polycistronic expression refers to the situation in which two (bicistronic) or more ORFs are encoded by a single messenger RNA (mRNA). Although historically considered incompatible with the mechanism of translation in eukaryotes, discoveries of nucleus-encoded loci that produce more than one protein have challenged this dogma. Polycistrons in which mature mRNAs contain two or more protein-coding ORFs have been detected in the transcriptomes of fungi (9), plants (10, 11), fruit flies (12, 13), and mammals (14). In many of these instances, the candidate loci are not exclusively polycistronic, with a population of monocistronic transcripts for each ORF also produced, which has raised questions about their importance. Moreover, the mechanism by which the downstream ORFs in polycistronic mRNAs are translated remains unknown in most of these cases.

Eukaryotic viruses have evolved unique mechanisms of non-canonical eukaryotic translation for expressing multiple polypeptides from a single mRNA. Internal Ribosome Entry Sites (IRES) in mammalian viruses contain distinct secondary structures that directly recruit translation initiation components to an ORF, independent of the 5’ cap (15). 2A “self-cleaving” peptide sequences contain motifs that trigger ribosomal skipping events, generating multiple polypeptides from a single ORF due to failure to form a peptide bond (16). Polycistronic expression is an attractive tool for coexpression of transgenes, and both IRES and 2A peptides have been widely used in expression vectors for producing multiple polypeptides (17, 18).

In eukaryotic genomes, variations from canonical translation can drive downstream ORF expression. In cases where two ORFs are in close proximity, ribosome components can reassemble and initiate translation of a downstream ORF following termination at an upstream ORF, a process termed post-termination reinitiation (19–21). In leaky ribosomal scanning, a suboptimal context sequence surrounding the initiator codon allows the scanning complex to occasionally bypass the first ORF, enabling translation of an alternative downstream ORF (21–23). These two competing models for translation of polycistronic ORFs can be distinguished by the role that is played by the sequence surrounding the initiator codon, referred to as the Kozak-like sequence. In the post-termination reinitiation model, a more favorable Kozak-like sequence of the most upstream ORF will increase translation of both ORFs. In the leaky ribosome-scanning model by contrast, a more favorable Kozak-like sequence of the upstream ORF will increase its translation at the expense of a downstream ORF.

Green algae (Chlorophyta) are a diverse clade of eukaryotes within the plant lineage (24). Like plants, they are primary producers that contribute significantly to carbon capture. Moreover, a few species serve as invaluable reference organisms for understanding metabolic processes such as photosynthesis (25) and are promising platforms for synthetic biology applications (26). We previously used long-read RNA sequencing (i.e. IsoSeq) to discover widespread and highly conserved loci encoding exclusively polycistronic mature mRNA in diverse green algae (27). Polycistronic genes contain ORFs that are independently translated, containing their own start and stop codons separated by a small spacer sequence (referred to as the “inter-ORF” region). Subsequent sequence analysis and proteomics confirmed the authenticity and translation of multiple ORFs at these loci in the algae, and raised the question of how the downstream ORFs are translated when they occur only in a polycistronic context. Polycistronic ORFs in the green algae vary in length and are often not read in the same frame. These features are incompatible with mechanisms such as 2A peptide motifs or stop codon read through, and there is no evidence that the inter- ORFs have any IRES-like activity (27). Additionally, in vitro transcription and translation of bicistronic loci from *Chlamydomonas reinhardtii* and *Chromochloris zofingiensis* revealed that the ratio of polypeptides produced from each ORF was dependent on the quality of the upstream (ORF 1) Kozak-like sequence (27). This observation suggests that translation of the downstream ORF (ORF 2) may depend on leaky scanning of the ORF 1 initiator codon, but this was not tested in vivo. In this work, we utilized two divergent (∼700 mya) green algae species, the classic reference organism *Chlamydomonas reinhardtii* and the emerging model alga *Auxenochlorella protothecoides,* to establish a high confidence dataset of bicistronic loci. We then used bioinformatic and in vivo mutational analyses to test the leaky scanning hypothesis as a mechanism for translation of bicistronic ORFs.

## Results

### Establishing a high confidence dataset of *Chla. reinhardtii* polycistronic loci

Previous structural annotation improvements identified 87 loci within the *Chla. reinhardtii* genome that were exclusively polycistronic (27). We refined this set to identify a high-confidence subset using the following criteria: 1) evidence for translation of ORFs based on ribosome profiling (28), and 2) conservation of the ORFs between the reference strain genome (CC-4532) and two divergent field isolate genomes (CC- 1952 and CC-2931) (29). This produced a refined set of 35 high-confidence bicistronic loci that passed all criteria (Supplementary Dataset S01), 31 of which were identified previously (27). The four newly identified genes include a deeply conserved bicistronic locus originally described in land plants that encodes the CDC26 cell cycle regulator and the TTM3 phosphatase (11).

The predicted amino acid sequences of the ORFs were used to search for bicistronic orthologs in six other chlorophyte species: *Volvox carteri*, *Dunaleilla salina* and *Chromochloris zofingiensis* (all Chlorophyceae), *Coccomyxa subellipsoidea* (Trebouxiophyceae), and *Ostreococcus lucimarinus* and *Micromonas pusilla* (Mamiellophyceae). Preservation of the bicistronic arrangement was inferred when pairs of orthologous ORFs were colinear (neighboring ORFs on the same strand) in the queried genome (Figure 1A & Supplementary Dataset S02). Out of 35 loci, 20 were bicistronic in at least one of the six chlorophytes. Validation that 19 of these 20 colinear loci are bicistronic was provided by Iso-Seq reads (*Chro. zofingiensis* (30) and *D. salina* (31)) and/or expressed sequence tags (ESTs) (*V. carteri*, *D. salina*, and *Cocc. Subellipsoidea* (*27*)), confirming conservation of bicistronic organization across hundreds of millions of years of evolution.

**Figure 1.**
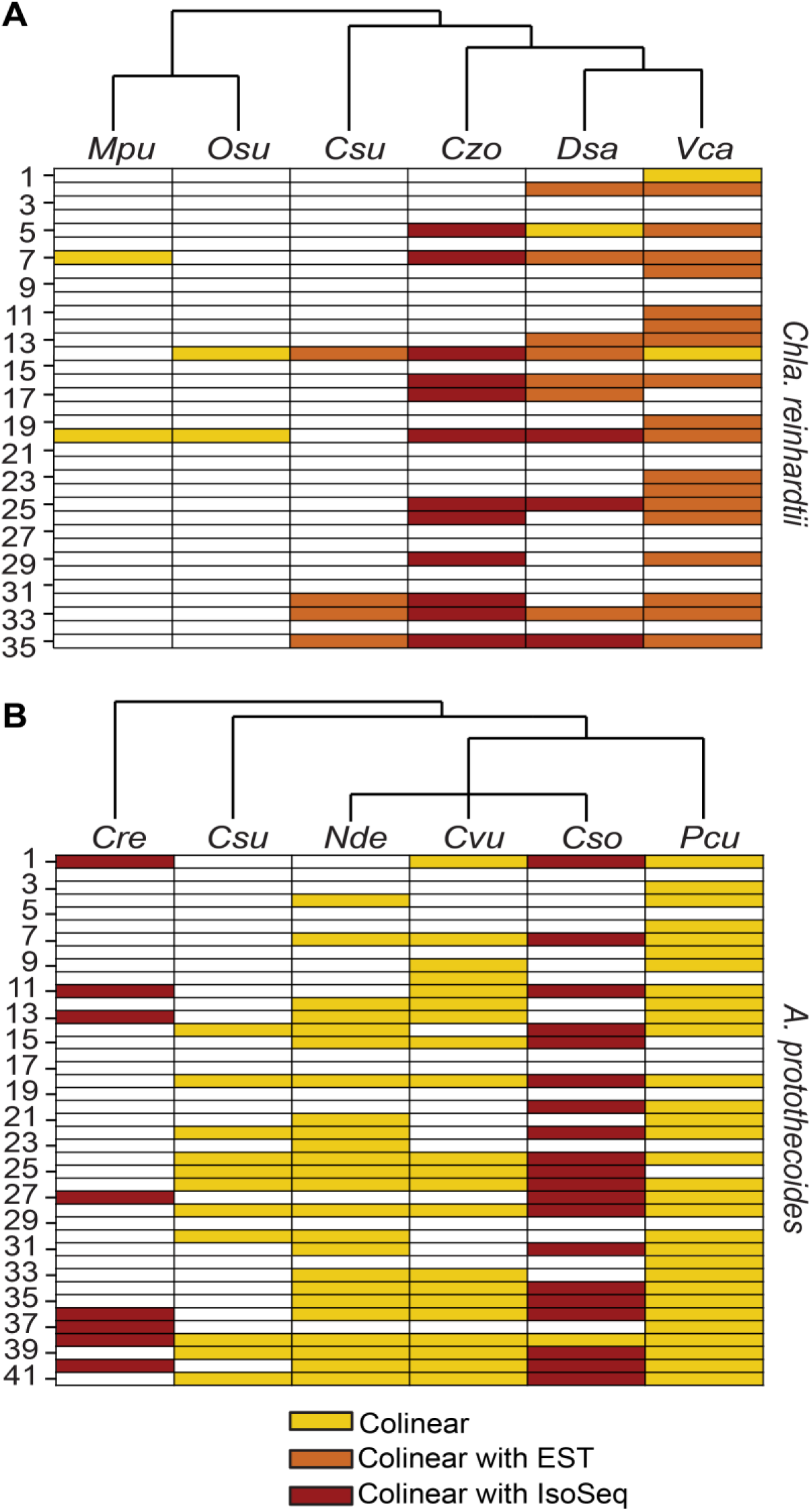
Bicistronic loci in divergent green algae. **(A)** 35 bicistronic loci in *Chla reinhardtii* are conserved in field isolates. Translation of both ORFs is supported by Ribo-Seq data. Homologs encoded in the bicistronic transcripts were identified in other Chlorophytes. High confidence hits and evidence of colinearity were further evaluated (See Methods). Yellow denotes a colinear pair of ORFs with significant similarity to a pair of bicistronic ORFs from *Chla. reinhardtii*, some of which were supported to be bicistronic with additional Iso-Seq or EST data, indicated in red and orange, respectively. Columns are ordered by the phylogenetic tree above each panel. Species are labeled according to the following code: *Cre, Chlamydomonas reinhardtii; Csu, Coccomyxa subellipsoidea; Czo, Chromochloris zofingiensis; Dsa, Dunaliella salina; Mpu, Micromonas. pusilla; Olu, Ostreococcus lucimarinus; Vca, Volvox carteri.* **(B)** 41 conserved and novel bicistronic loci were identified in *A. protothecoides*. A similar query search was performed to the one described in panel A but restricted within the Trebouxiophyceae with *Chla. reinhardtii* as an outgroup. Columns are ordered by the phylogenetic tree above each panel. Species are labeled according to the following code: *Pcu, Prototheca cutis; Cso, Chlorella. sorokiniana; Cvu, Chlorella vulgaris; Nde, Nannochloropsis desiccata; Csu, Coccomyxia subellipsoidea; Cre, Chlamydomonas reinhardtii*.

For reverse genetic analysis of polycistronic loci, we turned to the unicellular Trebouxiophyte alga *Auxenochlorella prototheocides*, which we have developed as a reference organism because of its small nuclear genome and experimental capability for genome manipulation by targeted homologous recombination (Craig et al, unpublished). Using Iso-Seq, we manually searched for loci in *A. protothecoides* where two or more ORFs were exclusively found on the same transcript. This uncovered 41 bicistronic loci within the genome of *A. protothecoides* strain UTEX 250-A (Supplementary Dataset S01), most of which had not been previously identified as polycistronic in other species.

The criteria described above were used to assess whether the bicistronic arrangement was conserved in other members of the Trebouxiophyceae: *Coccomyxa subellipsoidea*, *Chlorella vulgaris*, *Chlorella sorokiniana*, *Nannochloris dessicata*, and *Prototheca cutis*; and in *Chla. reinhardtii* (Figure 1B & Supplementary Dataset S02). 35 out of the 44 bicistronic loci in *A. protothecoides* had colinear hits in at least one other Trebouxiophyte, suggesting that they function as such. 22 of these loci were further validated by Iso-Seq reads in the transcriptomes of either *Chlo. sorokiniana* (32) or *Chla. reinhardtii* (27). Notably, eight loci are bicistronic in both *Chla. reinhardtii* and *A. protothecoides,* indicative of conservation over 700 million years of evolution (33). The set of 35 bicistronic loci from *Chla. reinhardtii* and 41 from *A. protothecoides* represent the high confidence set for downstream bioinformatic analyses.

### Structural features of green algal bicistronic genes

We next examined the structural properties of the curated sets of bicistronic loci to identify features differentiating them from monocistronic genes, reasoning that such features might give insight into the mechanism for translating multiple ORFs. In *Chla. reinhardtii*, bicistronic ORF 1 sizes were significantly smaller (median = 357 nt) than those of ORF 2 (median = 1113 nt) and monocistronic ORFs (median = 1,509 nt). ORF 1 sequences in the high confidence set were also shorter than previously described in the unrefined set (31) (median = 600 nt) (Figure S1B). The ORF size differential was also present in *A. protothecoides*, with ORF 1 (median = 291 nt) being significantly shorter than both ORF 2 (median = 1185 nt) and monocistronic ORFs (median = 1005 nt) (Figure S1E).

We also compared the distribution of the inter-ORF lengths for colinear genes (defined here as genes on the same chromosome strand with ≤20,000-nt separation between ORFs), and for predicted uORFs (small ORFs found in the 5′ untranslated regions of monocistronic genes). Consistent with our previous study (27), the inter-ORF spacing at bicistronic loci is much shorter than the distance between monocistronic colinear genes, both in *Chla. reinhardtii* (bicistronic median = 183 nt, monocistronic median = 3415 nt,) and *A. protothecoides* (bicistronic median = 40 nt, monocistronic median = 2518 nt,) (Figure S1C & S1F). The inter-ORF spacing of bicistronic genes from the refined *Chla. reinhardtii* and *A. protothecoides* sets were not statistically different from uORF-ORF spacing in either genome (median = 148 nt for *Chla. reinhardtii* and median = 76 nt for *A. protothecoides*) (Figure S1C & S1F).

Short 5’ UTRs or initiator codons very close to the 5’ cap have been associated with a higher degree of leaky scanning (34, 35). Bicistronic mRNA transcripts had significantly shorter 5’ UTRs (median = 83 nt for *Chla. reinhardtii*, and median = 54 nt for *A. protothecoides*) than monocistronic ORFs (median = 214 nt for *Chla. reinhardtii*, and median = 112 nt for *A. protothecoides*), which may contribute to more frequent leaky ribosomal scanning to bypass the ORF 1 initiation site for downstream ORF translation (Figure S1A & S1B).

### ORF 1 of bicistronic transcripts has a weak Kozak-like sequence

The Kozak-like sequence determines the recognition efficiency of a translation initiation site. In the context of leaky scanning, for translation of ORF 2 to occur, the Kozak-like sequence of ORF 1 should be suboptimal, allowing for the pre-initiation complex to occasionally scan through ORF 1. To address this, Kozak-like consensus sequences were deduced for both *Chla. reinhardtii* and *A. protothecoides* (Figure 2A and 2B). The Kozak-like consensus sequence varies between different eukaryotic lineages (36, 37), but has been defined broadly as gcc(A/G)ccAUGG (3), with A or G at the −3 position and G at the +1 position relative to the initiator codon as the most conserved elements. Indeed, both algae retain these conserved features (Figure 2A and 2B), but the *A. protothecoides* consensus is more GC-biased than that of *Chla. reinhardtii*, particularly at the + 1 position relative to the initiator codon.

**Figure 2.**
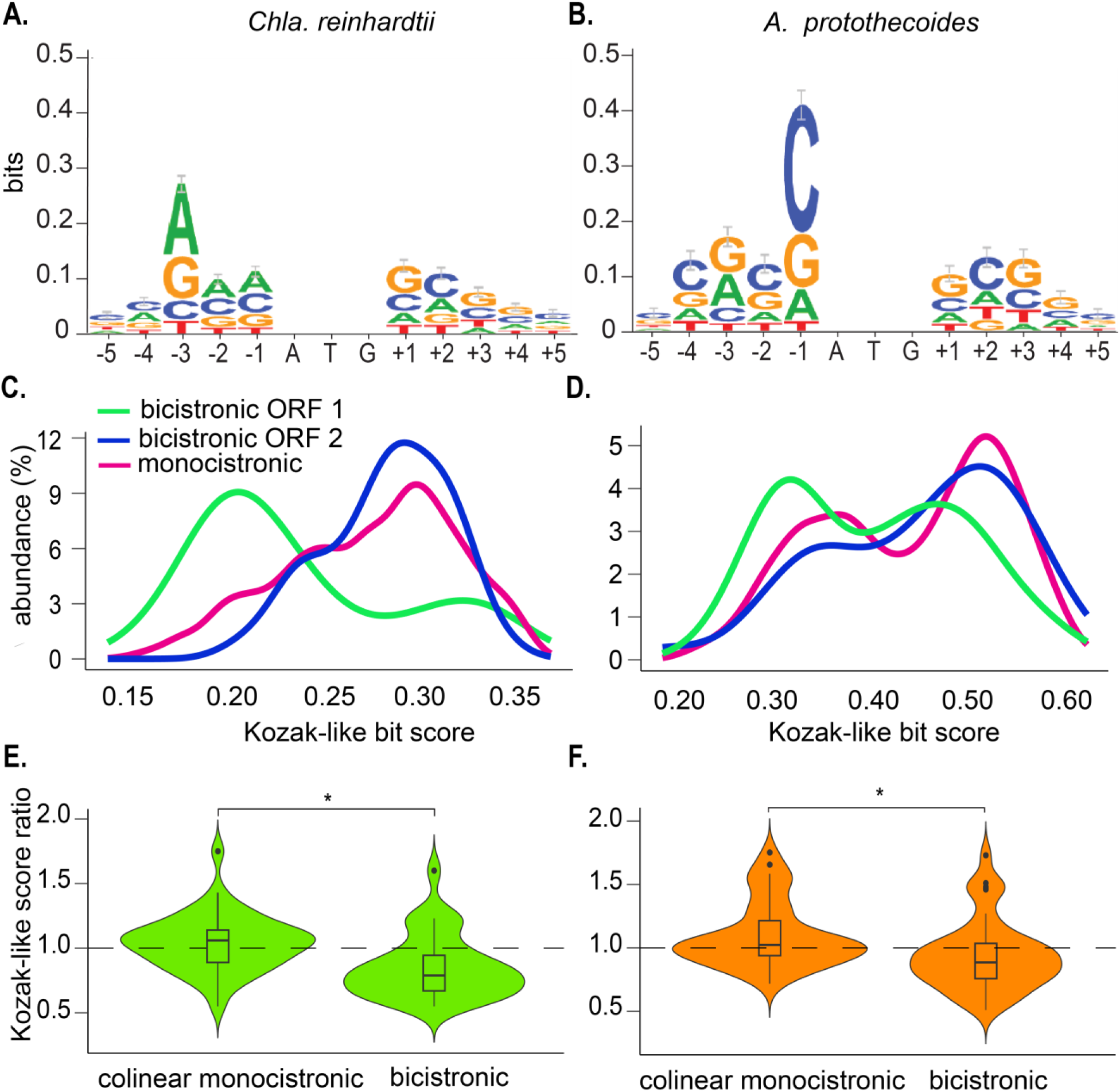
ORF 1 in bicistronic loci is associated with weak Kozak-like sequences. **(A)** Kozak-like consensus sequence for *Chlamydomonas reinhardtii*. The consensus sequence was generated by obtaining the surrounding context sequence of all initiation codons on monocistronic genes, of which a random subset (n=7210) was used to generate the WebLogo. (**B)** Kozak-like consensus sequence for *A. protothecoides* generated with a random subset of monocistronic genes (n=3824) The WebLogos in A and B were used to generate bit scores based on nucleotide bias at each position. (C) Scores were plotted for *Chla. reinhardtii* polycistronic ORF 1 (n=35), polycistronic ORF 2 (n=35), and the other half of the monocistronic genes from the genome not used in the initial WebLogo (n=3824). **(D)** Density plot of Kozak-like sequence bit scores for *A. protothecoides*. Values include polycistronic ORF 1 (n=41), ORF 2 (n=41), and monocistronic genes not used in generation of the WebLogo (n=3849). (**E and F)** Kozak-like score ratios in *Chla. reinhardtii* and *A. protothecoides* A ratio of the bit score of ORF 1 to ORF 2 was generated for all bicistronic genes (*n*=35 and n = 41 respectively) as a measure of transcript specific comparison. As a control, a random sample set (n=40) of colinear monocistronic genes (adjacent genes on the same strand and chromosome within ≤ 5,000 region between ORFs) were also compared. Dashed line denotes a score ratio of 1. Whiskers indicate 1.5 times the interquartile range, and notches indicate the confidence interval of the median. Outliers are plotted as individual points. Statistical significance was tested for with a Kruskal-Wallis test and Wilcoxon rank-sum test for sample comparison. Asterisks “*” indicated p-values less than .05.

We then used the consensus sequence to quantify the Kozak-like sequence strength of individual ORFs. WebLogo density bit scores indicate how well each Kozak-like sequence resembles the deduced consensus (presumed optimal) sequence (38, 39). Bit scores were calculated for monocistronic and bicistronic genes based on five nucleotides upstream and downstream of the annotated initiation codons (Figure 2C). For each species, half of the monocistronic genes, chosen at random, were used to produce the original consensus sequence, while the remaining half were used to calculate the distribution of bit scores. In *Chla. reinhardtii*, the distribution of scores for ORF 1, ORF 2, and monocistronic genes reveals a large proportion of weak scores for bicistronic ORF 1, in contrast to the significantly higher scores of bicistronic ORF 2 and the monocistronic ORFs (Figure 2C). This trend, although less pronounced, is also observed in *A. protothecoides* (Figure 2D). These distributions are in agreement with ORF 1 of bicistronic loci being permeable to intermittent leaky scanning. In contrast to ORF 1, ORF 2 would be expected to have a stronger Kozak-like sequence to maximize its translation. We tested this assumption by comparing the ratios of ORF 1 to ORF 2 Kozak-like bit scores for each individual bicistronic locus with the ratios of a random selected set of neighboring monocistronic genes (within 3,000 bp of each other). Given the propensity for a weak Kozak-like sequence at ORF 1 and a stronger Kozak-like sequence at ORF 2, we expected ratios of less than 1 for the bicistronic ORFs. Conversely, since mRNA transcripts from adjacent genes are translated independently, the Kozak-like sequence of one transcript should have no influence on the other, and we expected ratios close to 1 for the monocistronic genes. In both organisms, the average ratio for the bicistronic genes was significantly smaller (median = 0.79 for *Chla. reinhardtii*, median = 0.88 for *A. protothecoides*) than for the monocistronic control sets (median = 1.06 for *Chla. reinhardtii*, median = 1.02 for *A. protothecoides*), indicating that the Kozak-like sequences of each ORF are in a favorable configuration for leaky ribosome scanning to occur (Figure 2E & 2F).

### Bicistronic genes show start codon depletion in regions before ORF 2

The occurrence of start codons upstream of the authentic initiation codon can inhibit translation of a downstream ORF (40–42). To test the frequency of alternative initiation codons in bicistronic loci, we calculated the observed frequency of “ATG” 3-mers within regions of the bicistronic loci and compared it to an expected estimation based on the sequence length (see Methods & Supplementary Dataset S02). The probability was adjusted based on the GC sequence bias of the *A. protothecoides* and *Chla. reinhardtii* transcriptomes to calculate the expected frequency of “ATG” sequences in the 5’ UTR, ORF 1, inter-ORF, the total cumulative sequence upstream of ORF 2, and ORF 2 for all bicistronic loci, and in monocistronic loci for comparison.

To assess the validity of this method for predicting the frequency of start codons, we calculated Pearson Correlation Coefficients (PCCs) between “ATG” frequencies within 5’ UTRs and CDS, and their sequence lengths for sets of monocistronic genes. There was a high positive correlation between the “ATG” count and the 5’ UTR plus CDS sequence length (*r* = .89 for *Chla. reinhardtii*, and *r* = .76 for *A. protothecoides*) for both algae, indicating that this was a valid metric for predicting “ATG” sequence bias (Figure S2). Observed/expected ratios were compared for ORF 2 and regions upstream of ORF 2 in bicistronic transcripts (Figure 3A and 3B). In theory, a sequence with no bias against “ATG” sequences should have an observed/expected ratio value of 1. Values less than 1 indicate bias against “ATG” sequences. The observed/expected values of the 5’ UTRs in monocistronic genes had median values of .47 and 0 in *Chla. reinhardtii* and *A. protothecoides* respectively. Assuming correct annotation of the initiation codons, this bias against ATG sequences in the 5’ UTRs is consistent with maximizing ORF translation. Median observed/expected values for the monocistronic CDS and for the combined 5’ UTR and CDS sequences ranged from 0.94-1.20 in both algae (Figure 4A and 4B), suggesting no discrimination against ATG 3-mers in these regions. The observed/expected ratios for bicistronic ORF 2 had similar values, with medians of 0.99 and 1.20 for *Chla. reinhardtii* and *A. protothecoides* respectively. However, we observed a striking reduction in observed/expected ratios in the regions upstream of bicistronic ORF 2 for both algal species, with medians values of .49 and .43 for the sum of the sequence before ORF 2 in *Chla. reinhardtii* and *A. protothecoides* respectively. (Figure 3A and 3B). This bias against potential alternative start sites is consistent with optimization for ribosome scanning to ensure correct translation of ORF 2. The relatively small size of bicistronic ORF 1 regions (Figure S1B, S1E) may help to minimize the frequency of translation from alternative initiation sites.

**Figure 3.**
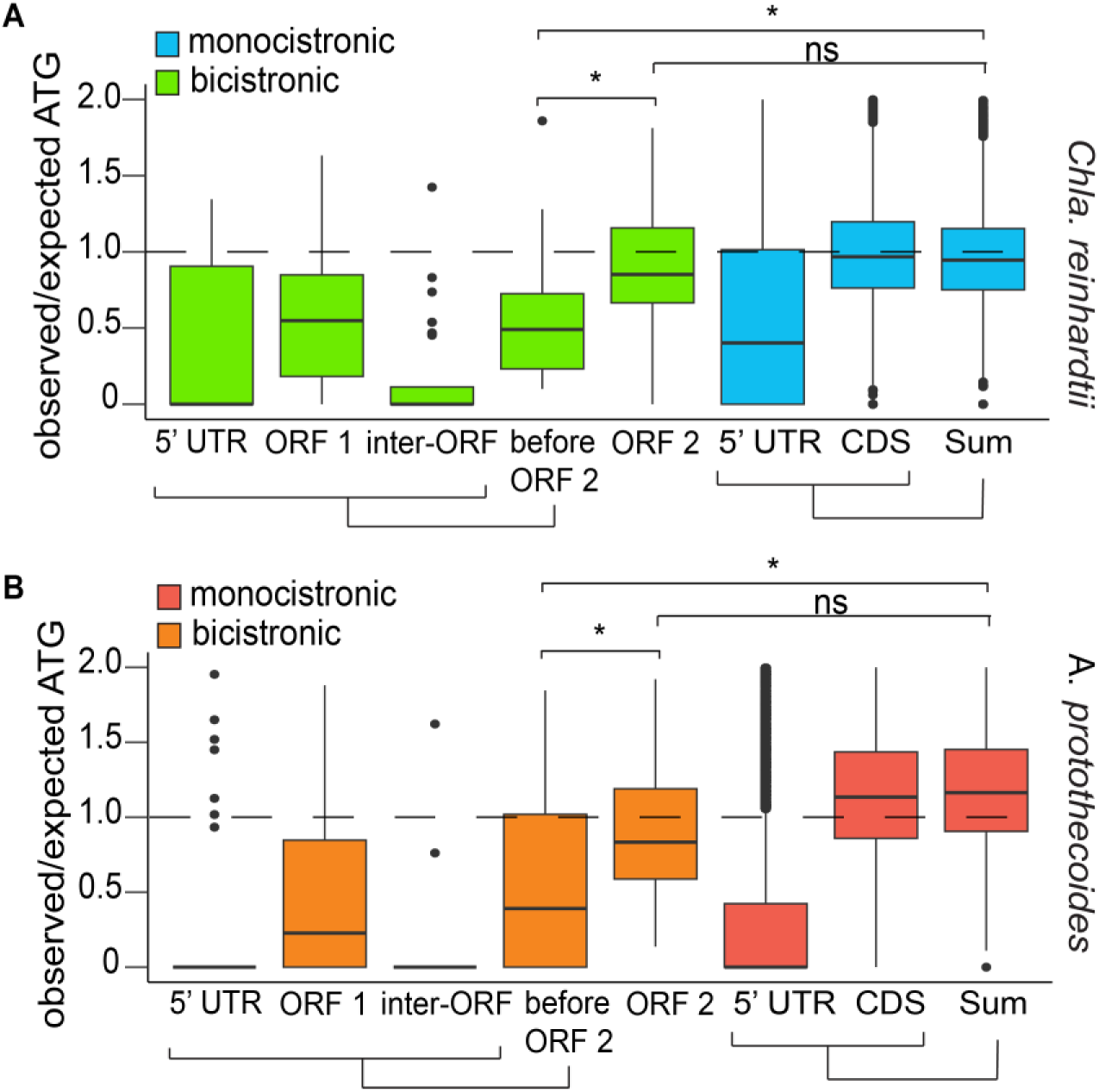
A bias against start codons upstream of ORF 2 in bicistronic loci. **(A and B)** Distribution of observed over expected ratios of “ATG” sequences based on nucleotide length. The presence of an ATG was estimated based on sequence length and the random chance of encountering an ATG in the reading strand. For the bicistronic (n= 35 & n = 41) set, this ratio was calculated for both individual components of the transcripts (“5’ UTR, “ORF 1”, “Inter-ORF”, “ORF 2”) and the sum of the components before the start codon of the downstream ORF (“before ORF 2”). For monocistronic genes (n=15333 & n=7698), the analysis was done for all annotated 5’ UTR, CDS, and the sum of those regions (Sum). Whiskers indicate 1.5 times the interquartile range, and notches indicate the confidence interval of the median. Outliers are plotted as individual points. Statistical significance was tested for with a Kruskal-Wallis test and Wilcoxon rank-sum test for sample comparison. Asterisks “*” indicated p-values less than .0001.

**Figure 4.**
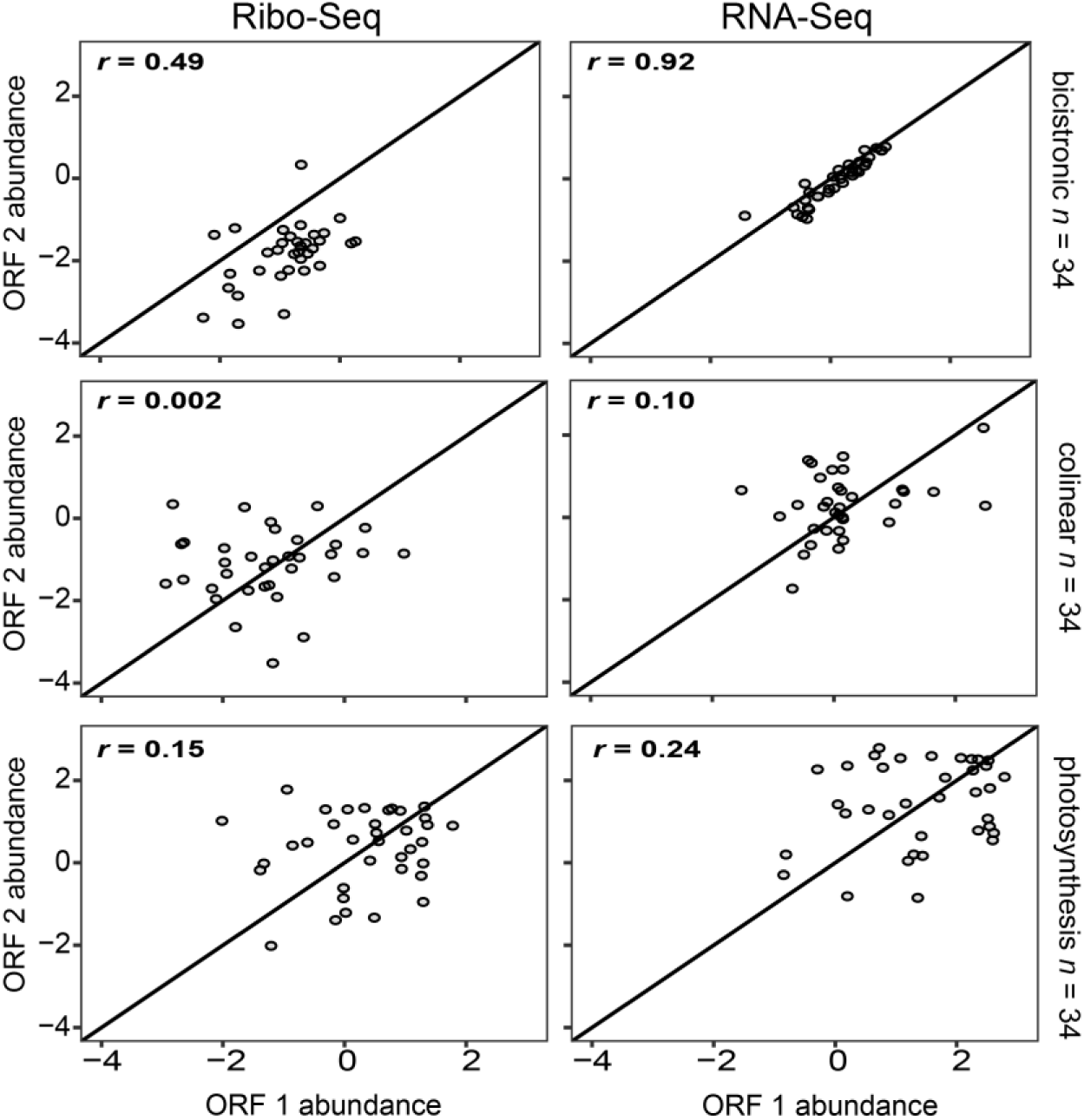
Correlation of RNA and ribosome occupancy in *Chlamydomonas reinhardtii*. Correlation analysis of RNA-Seq reads and Ribo-Seq reads for pairs of bicistronically expressed genes (n = 34). Log_10_ -normalized coverage scores (counts) per nt of transcript) were plotted with ORF 1 (upstream) on the x-axis and ORF 2 (downstream) on the y-axis. A Pearsons correlation coefficient (*r*) was calculated as indicated. A diagonal line representing a perfect 1:1 correspondence is plotted for reference. The same analysis was applied to monocistronically expressed, colinear genes which are defined here as adjacent genes on the same strand of the same chromosome with ≤10,000 nt between ORFs. Random sampling of 34 colinear gene pairs and their *r* values are presented here. Lastly, pairs of genes annotated by Gene Ontology to be in a common pathway (GO:0015979 photosynthesis) were paired randomly and analyzed as above (n = 36).

### Correlation of RNA and ribosome abundance in *Chla. reinhardtii*

To better understand how bicistronic loci behave in vivo, we first turned to genome-wide expression patterns in the transcriptome and translatome of *Chla. reinhardtii*. Previous analyses have demonstrated that authentic polycistronic transcripts display nearly identical mRNA abundance estimates for each individual ORF (9, 27). To test this for our filtered sets of bicistronic genes, we calculated PCC values for the bicistronic ORFs using log10 normalized reads of pooled transcriptomic datasets (27). We further extended this analysis to log10 normalized reads from a ribosome profiling dataset (28) to compare ribosome occupancy at each ORF. For comparison, two control sets were assembled, one consisting of a random sample of colinear genes, and another set of randomly paired photosynthesis-related genes, as predicted by gene ontology (GO) annotation (43). The colinear genes showed correlation coefficient (*r*) values of 0.10 and 0.002 for the RNA-Seq and Ribo-Seq, indicative of negligible correlation as expected (Figure 4). To ensure that this result did not occur from sampling bias, we repeated this analysis for the whole population of colinear genes, which showed a similar lack of correlation (Figure S3). Although higher than what was seen in colinear genes, photosynthetic gene pairs still showed negligible correlation of read abundance and ribosome occupancy (Figure 4, *r* =0.24 and *r* = 0.15 for RNA-Seq and Ribo-Seq respectively). As we observed previously (27), there is a very high positive correlation between ORF 1 and ORF 2 RNA-seq read abundances in the bicistronic dataset (*r* = 0.92, Figure 4, top right). While not as highly correlated as the RNA-seq abundances, a low positive correlation was observed in ribosome occupancy between ORF 1 and ORF 2 (*r* = 0.49, Figure 4, top left). Additionally, Ribo-seq read abundance was almost always higher for ORF 1 than for ORF 2 (32/35 loci) (Figure 4, top left), implying lower levels of ORF 2 translation. This trend is consistent with observed Ribo-seq patterns in viral polycistrons in which alternative translation initiation events such as leaky scanning drive multiple ORFs (44), and with protein abundance patterns in experimentally analyzed bicistronic loci in other eukaryotes (45, 46).

### In vivo manipulation of ORF 1 Kozak-like sequence influences expression of ORF 2

Having established that the structural patterns, sequence properties, and expression correlations of bicistronic loci were compatible with the leaky scanning mechanism, we sought to further test the mechanism *in vivo*. For these experiments, high frequency gene targeting by homologous recombination in *A. protothecoides* was utilized to allow transgene insertions and mutational analyses with minimal positional effects. Quantifying the expression of endogenous bicistronic genes, many of which have unknown functions, posed a challenge; consequently, we designed a reporter system by cloning endogenous bicistronic loci and replacing ORF 2 with the CDS of *Venus*. These designs were applied to two bicistronic loci: *TOM22_SDHAF3* (UTEX250_A10.30135/UTEX250_A10.30130) and *OST4_FAM32A* (UTEX250_A05.11130/UTEX250_A05.11130) (Figure S4), selected on the basis of adequate mRNA expression for Venus detection, and conservation between *A. protothecoides* and *Chla. reinhardtii* (supplementary dataset 1, S01). These modified bicistronic loci were included in a gene cassette that also contained a selectable marker module conferring G418 antibiotic resistance and were integrated into the UTEX 250-A genome at a neutral homozygous landing site (*DAO1*) using targeting flanks (Figure 5A).

**Figure 5.**
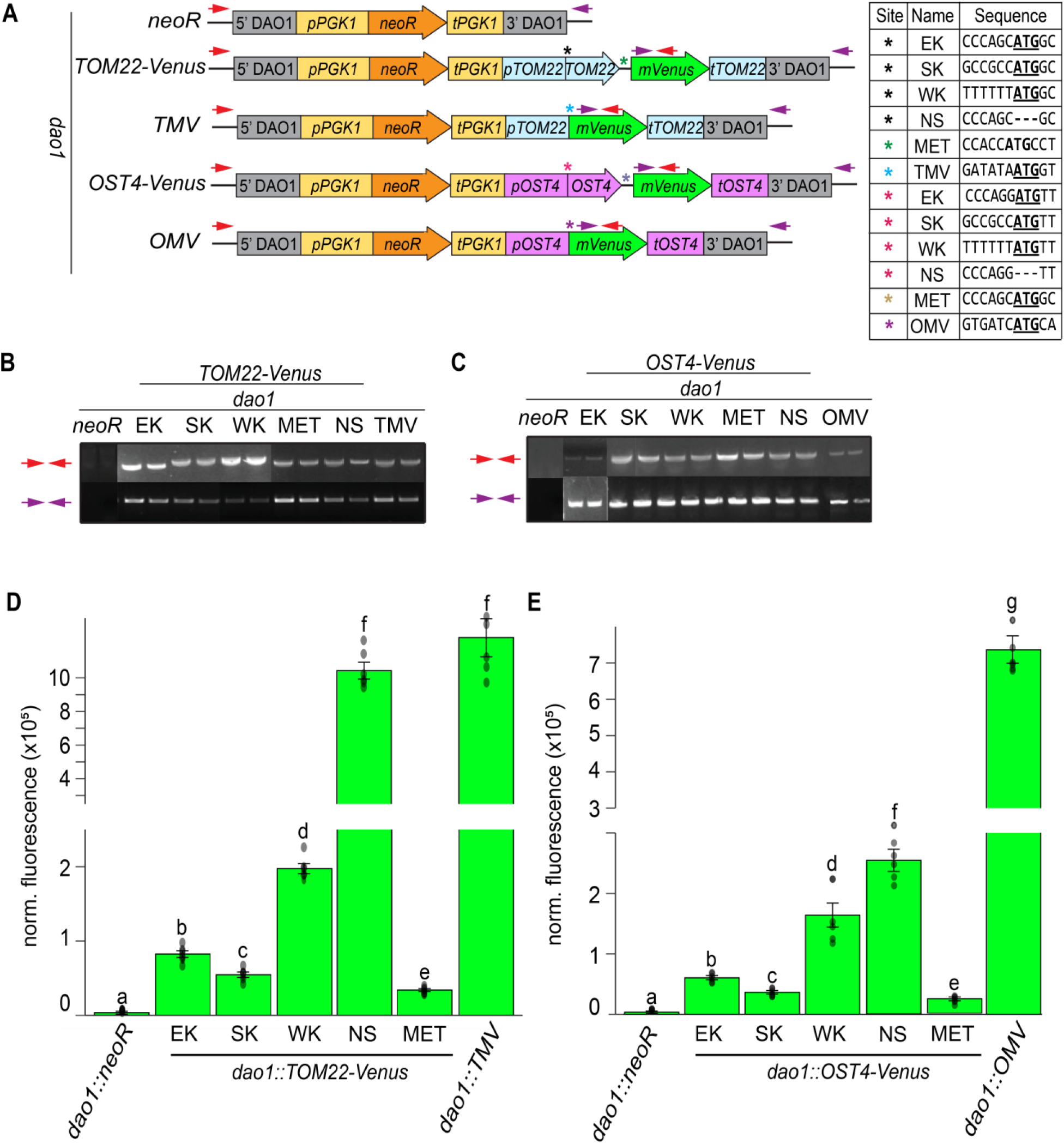
The Kozak-like sequence and start codon of ORF 1 influences ORF 2 expression in vivo. **(A)** Construct designs for targeted integration of a Venus reporter module into the *DAO1* neutral locus in *A. protothecoides*. Targeting 5’ and 3’ flanks are depicted in light grey. For G418 selection, a *neoR* module consisting of a promoter sequence from *ApPGK1* (*pPGK1*), the NeoR CDS (*NeoR*), and a terminator sequence from *ApPGK1* (*tPGK1*) is indicated in yellow and orange. Blue *TOM22* and pink *OST4* labels indicate endogenous components of the two individual polycistronic loci respectively. *p* and *t* denote the promoter and terminator regions respectively. ORF 2 was swapped with the CDS of mVenus depicted in lime. The inter-ORF is denoted by the straight black before the *mVenus*. Asterisks denote sites of mutational analysis, with specific sequences shown in the table. EK= Endogenous Kozak, SK = Strong Kozak, WK = Weak Kozak, NS = No start, TMV = TOM9 Monocistronic Venus, OMV = OST4 Monocistronic Venus. MET denotes a construct with the endogenous ORF 1 Kozak-like sequence but an additional out of frame ATG in the inter-ORF. Initiator codons are bolded and underlined. Nucleotide deletions are denoted as dashes. Red and purple arrows denote primers. **(B and C)** Genomic PCR amplification of the TOM22-Venus and Ost4-Venus cassettes at the *DAO1* locus. Red and purple primer pairs correspond to regions shown in panel A. For the Tom22-Venus, red primers bands correspond to a size of 4681 bp for EK, SK, and WK, 4684 bp for MET, 4681 bp for NS, and 4192 bp for TMV. *neoR* displays no band. Purple primers bands are 1908bp for all strains except *neoR*, which displays no band. 2 Representative bands are shown for each strain. For OST4-Venus, red primers bands correspond to a size of 3966 bp for EK, SK, and WK, 3969 bp for MET, 3963 bp for NS, and 3765 bp for OMV. NeoR displays no band. Purple primers denote bands that are 1908bp for all strains except NeoR, which displays no band. 2 representative bands are shown for each strain. **(D and E)** Normalized relative fluorescence output of downstream Venus test for the Tom22-Venus and OST4-Venus constructs. For both D and E, statistical significance was tested for with the Kruskal-Wallis test and Wilcoxon rank-sum test for sample comparison. Letters indicate groups of comparative statistical significance with p-values less than .05.

Variant constructs were designed in which the endogenous Kozak-like sequence (EK) of ORF 1 was mutated to be either stronger (SK) or weaker (WK) based on optimal and suboptimal Kozak-like sequences (Figure 3B). Mutations were only introduced upstream of the initiation codon of ORF 1 to avoid altering the protein coding sequence (Figure 5A). To test whether the presence of ORF 1 influences ORF 2 expression, a construct was created in which the ORF 1 start codon was removed entirely (NS). To evaluate the influence of upstream alternative initiation codons on *Venus* translation, an additional construct was designed that preserved the endogenous ORF 1 Kozak-like sequence but contained an additional initiation codon in the inter-ORF region out of frame with the *Venus* CDS (MET). A construct that only contains the selectable marker module (*neoR*) targeted to *DAO1* provided a negative control, and monocistronic *Venus* driven by the endogenous promoters with the ORF 2 Kozak-like sequence (TMV/OMV) provided a positive control.

All strains were genotyped via PCR amplification of the gene cassette and neighboring sequence to ensure proper integration at the *DAO1* neutral landing site prior to phenotypic analysis (Figure 5A and 5B). Although precise integration of the constructs should minimize variability of reporter expression due to position effects, we measured the abundance of the bicistronic transcripts encoding *Venus* using Real-Time qPCR to test for significant differences in transcription or mRNA maintenance among the strain*s*. In both the *TOM22-Venus* and *OST4-Venus* contexts, there was no significant difference in transcript accumulation between strains (Figure S5), suggesting that differences in Venus fluorescence between strains should be attributed to *Venus* translation rather than template abundance.

Venus fluorescence was substantially higher in both *dao1::TOM22-Venus* and *dao1::OST4-Venus* strains than the background fluorescence detected in selectable marker-only (*dao1::neoR*) negative control lines (Figure 5D and 5E), but all bicistronic configurations showed significantly less fluorescence than the monocistronic Venus positive controls (*dao1::OMV*, *dao1::TMV*). A notable exception was the *dao1::TOM22-Venus* strain lacking the *TOM22* start codon (NS), which had almost as much fluorescence signal as the positive control (Figure 5D). Similarly, removal of the ORF 1 initiator codon (NS) from *dao1::OST4-Venus* spiked *Venus* fluorescence over 5-fold in comparison to strains in which the integrity of ORF 1 is maintained (Figure 5E). These results suggested that translation of ORF 1 inhibited translation of the downstream *Venus* ORF. Next, we examined the effects of altering the ORF 1 Kozak-like sequence on *Venus* expression, hypothesizing that an inverse relationship between the strength of the ORF 1 Kozak-like sequence and Venus fluorescence would be consistent with a leaky scanning mechanism. Fluorescence decreased by factors of 1.5 and 1.7, respectively, in strains expressing *dao1::TOM22-Venus* and *dao1::OST4-Venus* with a stronger ORF 1 Kozak-like sequence (SK) compared to the endogenous Kozak-like sequence (EK) (Figure 5D, 5E). In contrast, weaker Kozak-like sequences (WK) increased fluorescence over 2-fold in the context of both *dao1::TOM22-Venus* and *dao1::OST4-Venus* (Figure 5D and 5E). Introduction of an out of frame initiation codon into the inter-ORF regions (MET) led to a significant decrease in Venus fluorescence compared to all other strains, with less than half of the expression of EK strains. These results are consistent with alternative potential translation start sites limiting ORF 2 expression (Figure 5D and 5E).

### A synthetic bicistronic locus displays coexpression and tunability via the Kozak-like sequence

In light of the observation that manipulating ORF 1 Kozak-like sequences influences ORF 2 translation in vivo, we designed a synthetic bicistronic dual reporter to quantify the coexpression of multiple transgenes and evaluate the effect of Kozak-like sequence manipulation. Our design employed a regulatable promoter to maximize signal output and incorporated an inter-ORF region from an endogenous bicistronic transcript. For high levels of transcription, we utilized the promoter and terminator regions from the gene encoding Photosystem I subunit D (*PSAD1)*, which in *A. protothecoides* is highly induced in photoautotrophy and repressed in heterotrophy. Consequently, we could leverage the reversible trophic switch to inhibit or induce expression (Figure 6A). Conveniently, the predicted size of the *PSAD1* 5’ UTR is also close in size (46 nt) to the median length for endogenous bicistronic transcripts (Figure S1). The *PSAD1* Kozak-like region is close to the consensus, maintaining the highly conserved elements at the −3 and −1 positions (Figure 6A).

**Figure 6.**
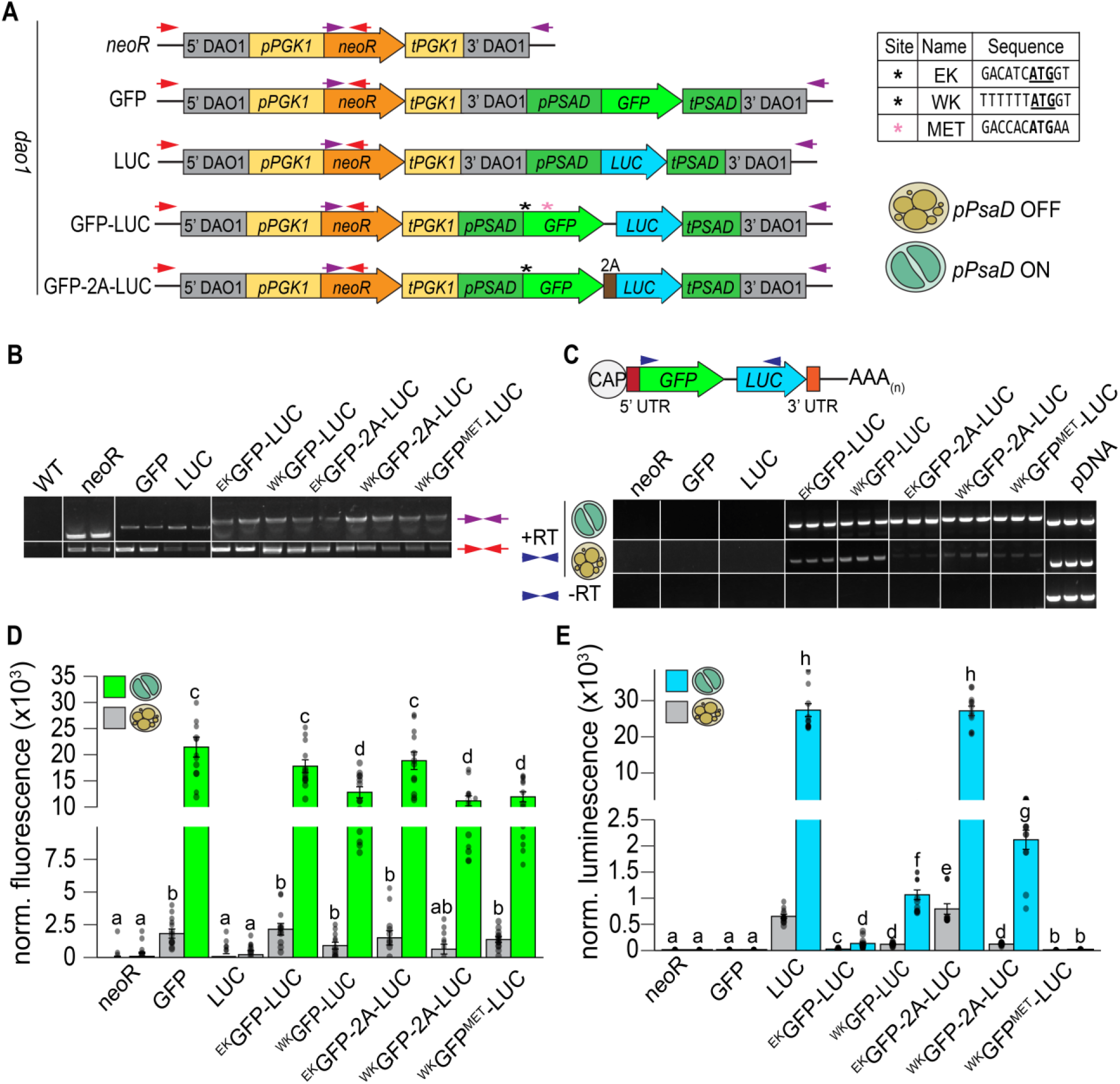
A synthetic bicistronic transcript displays tunable activity via the strength of the Kozak-like sequence. **(A**) Construct designs for a synthetic bicistronic dual reporter in *A. protothecoides*. *DAO1* targeting flanks and G418 resistance components are the same as previously described in figure 5A. The *REX1* inter-ORF is denoted as the black line before the Luciferase ORF. Bicistronic expression of the SuperFolder *GFP* (Lime) and *Gaussia princeps* Luciferase (*LUC*) denoted in blue were driven by the *PSAD1* promoter (dark green). Black asterisks denote sites of mutational analysis, with specific sequences shown in the table. EK= Endogenous Kozak, WK = Weak Kozak. MET denotes a construct with the reintroduction of an ATG codon into the CDS of the GFP (pink Atserisk). The *PSAD1* promoter is inducible dependent on either the phototrophic (green) or heterotrophic (yellow) state of *A. protothecoides*. **(B)** Genomic PCR amplification of all constructs at the DAO1 locus. 2 representative bands are shown for each strain. **(C)** Schematic of bicistronic RNA and cDNA blots of all transformed strains. Blue arrows denote primers for amplification. RT indicates the presence (+) or absence (-) of reverse transcriptase. Bands correspond to a size of 1,002 bp for strains producing the bicistronic transcript with the inter-ORF, and 1051 bp with for strains producing the bicistronic transcript with a 2A sequence. 3 representative bands are shown for each strain. **(D)** Normalized GFP fluorescence (488/528 ex-em) of all transformed strains with the *PSAD1* promoter off (gray) and on (green). Readings are normalized according OD750 and the background fluorescence of wildtype UTEX 250 cultures. **(E)** Normalized luminescence (487nm) readings of all transformed strains with the *PSAD1* promoter off (gray) and on (blue). Readings are normalized according OD750 and the background fluorescence of wildtype UTEX 250 cultures. For D and E, statistical significance was tested for with the Kruskal-Wallis test and Wilcoxon rank-sum test for sample comparison. Letters indicate groups of comparative statistical significance with p-values less than .05.

In the ORF 1 position, we incorporated a SuperFolder *GFP* (47) codon optimized for UTEX 250-A, which was modified to replace five codons encoding methionine with leucine (48), ensuring that there are no potential in- or out-of-frame initiation codons within the CDS. In the downstream ORF 2 position, we placed a luciferase (*LUC*) CDS from *Gaussia princeps*, in which a signal peptide from *A. protothecoides* CAH1 was included for secretion (49). The ORFs were separated by the 14 bp inter-ORF sequence from the endogenous *REX1S-REX1B* bicistronic locus (UTEX250_A11.30960/ UTEX250_A11.30955, Craig et al, unpublished). We then tested the effects of maintaining (^EK^GFP-LUC) or weakening (^WK^GFP-LUC) the *PSAD1* Kozak-like sequence in bicistronic reporters. A bicistronic construct with a weak ORF 1 Kozak-like sequence and a single methionine codon reintroduced into the *GFP* CDS (^WK^GFP^MET^-LUC) was used to test the effect of alternative upstream translation initiation sites on ORF 2 expression. Positive control strains expressed monocistronic *GFP* or *LUC* driven by the *PSAD1* promoter with the endogenous Kozak-like sequence (Figure 6A). For comparison, we also included strains in which the stop codon of the GFP was removed and the inter-ORF was swapped with a 2A self-cleaving peptide sequence, so that *GFP* and *LUC* were translating a single ORF split into two polypeptides. This was conducted in the context of the endogenous (^EK^GFP-2A-LUC) and weak (^WK^GFP-2A-LUC) Kozak-like sequences, serving as positive controls for coexpression.

Transformed strains were genotyped via PCR amplification to ensure precise insertion at the *DAO1* landing site (Figure 5A, 5B, primers illustrated by red and purple arrows). Strains with correct insertions were used for further analysis. Transgene mRNAs were detected by RT-PCR in all lines using primers spanning the *GFP* and *LUC* ORFs (Figure 6C). Transcripts of the expected size were present in all strains (^EK^GFP-LUC, ^WK^GFP-LUC, ^EK^GFP-2A-LUC, and ^WK^GFP-2A-LUC, and ^WK^GFP^MET^-LUC), and were absent in negative (*neoR*) and monocistronic positive controls (GFP and LUC). RT-PCR products with *GFP* and *LUC* separated by either inter-ORF or 2A sequences were confirmed by sequencing (Figure S6). No amplification products were detected in controls run without reverse transcriptase, confirming the absence of gDNA contamination (Figure 6C). Transcript abundance was substantially reduced in heterotrophy compared to photoautotrophy in all strains, consistent with control of transgene mRNA accumulation by *PSAD1* regulatory sequences (Figure 6C).

RT-qPCR was conducted using ORF-specific primers for *GFP* and *LUC* to test for differences in RNA abundance in strains expressing the reporter constructs (Figure S7). No significant differences in transgene expression were observed between strains grown under photoautotrophic conditions, indicating that variations in GFP and luciferase activity were due to differential translation rather than transcript abundance. However, both GFP and luciferase reporter activities were considerably higher in photoautotrophic compared to heterotrophic cells (Figure 6D, 6E), consistent with increased transgene expression (Figure 6C). GFP fluorescence and luciferase activity were detected in all strains with bicistronic expression, except for ^WK^GFP^MET^-LUC, which had background levels of luciferase activity. This was likely a result of inefficient translation of the ORF 2 luciferase CDS due to the presence of an alternative initiation site within the *GFP* CDS (Figure 6E). Again, we examined the effects of manipulating the ORF 1 Kozak-like sequence on *ORF 2* translation. In all strains expressing *GFP* with a weak Kozak-like sequence (^WK^GFP), GFP fluorescence decreased significantly in comparison to those with the endogenous Kozak-like sequence (^EK^GFP) (Figure 6D). Conversely, luciferase activity increased nearly 10-fold in ^WK^GFP-LUC strains compared to ^EK^GFP-LUC strains (Figure 6E). In contrast, the activities of both reporters were positively correlated in response to Kozak-like sequence strength in strains expressing GFP and LUC as a single ORF, linked by a 2A peptide. GFP fluorescence was reduced by 2-fold and luminescence was reduced by 10-fold in ^WK^GFP-2A-LUC strains compared to monocistronic controls (GFP and LUC) and ^EK^GFP-2A-LUC strains, consistent with reduced translation of the GFP-2A-LUC ORF (Figure 6D, 6E). The more dramatic decrease in luciferase activity compared to GFP fluorescence in the ^WK^GFP-2A-LUC strain could be attributed to inefficient cleavage of the 2A peptide (52) impeding luciferase protein function or secretion. The 2A peptide sequence does not contain any alternative start sites or internal ORFs, hence both translation and processing of GFP-2A-LUC and leaky ribosome scanning could contribute to LUC translation; this may explain the higher levels of luciferase activity in ^WK^GFP-2A-LUC compared to ^WK^GFP-LUC strains (Figure 6E). GFP fluorescence and luciferase luminescence were visualized for further validation of expression (Figure S8).

## Discussion

### Evidence for Leaky Scanning as the mechanism for multiple ORF translation

In this study we have tested hypotheses to identify the primary mechanism for translation of multiple ORFs in exclusively bicistronic green algal transcripts. We find that the structural features of bicistronic genes are consistent with episodic leaky ribosome scanning for bypass of ORF 1 to translate ORF 2, and do not support ribosome reinitiation or IRES as significant contributors to translation of downstream ORFs. Bicistronic loci in our high-confidence *Chla. reinhardtii* and *A. protothecoides* sets have short 5’ UTRs (Figure S1A), weak Kozak-like sequences at the translation initiation site of ORF 1 (Figure 2), and display bias against alternative initiation sites upstream of ORF 2. (Figure 3). Similar observations were reported in an earlier study describing *stoned* and *Snapin*, two bicistronic loci in *Drosophila melanogaster* (50). The upstream ORFs at both Drosophila loci are deficient in internal AUG codons, and the Kozak-like regions diverged from the consensus (50). Consistent with our findings, here too, in vivo mutational analysis using bicistronic reporter constructs indicated that translation of the downstream ORF was dependent on leaky ribosome scanning (50).

ORF 1 and ORF 2 Ribo-seq reads in bicistronic Chlamydomonas mRNAs were only moderately correlated, whereas there was a very high positive correlation between the RNA-seq reads. Reduced ribosome occupancy in ORF 2 versus ORF 1 is consistent with a mechanism in which there is independent translation of each ORF (Figure 4). Further in vivo studies of bicistrons demonstrated ORF 2 translation is inversely correlated with the strength of the Kozak-like sequence of ORF 1, inconsistent with either an IRES-dependent mechanism or ribosome reinitiation. The downstream ORFs of two *A. protothecoides* bicistronic loci were replaced with *Venus*, and fluorescence could be turned up or down by manipulating the ORF 1 Kozak-like region (Figure 5). This in vivo outcome replicates the results of similar manipulations with *Chla. reinhardtii* and *Chr. zofingiensis* bicistronic mRNAs in vitro (27). A synthetic bicistronic gene cassette, designed in accordance with the structural features of endogenous bicistronic genes, successfully co-expressed the modified SuperFolder GFP and luciferase. Again, weakening the Kozak-like sequence of the first ORF (GFP), increased expression of the downstream luciferase, and conversely, luciferase activity was eliminated by the introduction of alternative initiation sites within the GFP CDS (Figure 6). Together, these results solidify a model in which green algal bicistronic expression occurs via cap-dependent initiation of ORF 1, coupled with episodic leaky ribosome scanning events that allow the pre-initiation complex to scan through ORF 1 and initiate ORF 2 translation (Figure 7).

**Figure 7.**
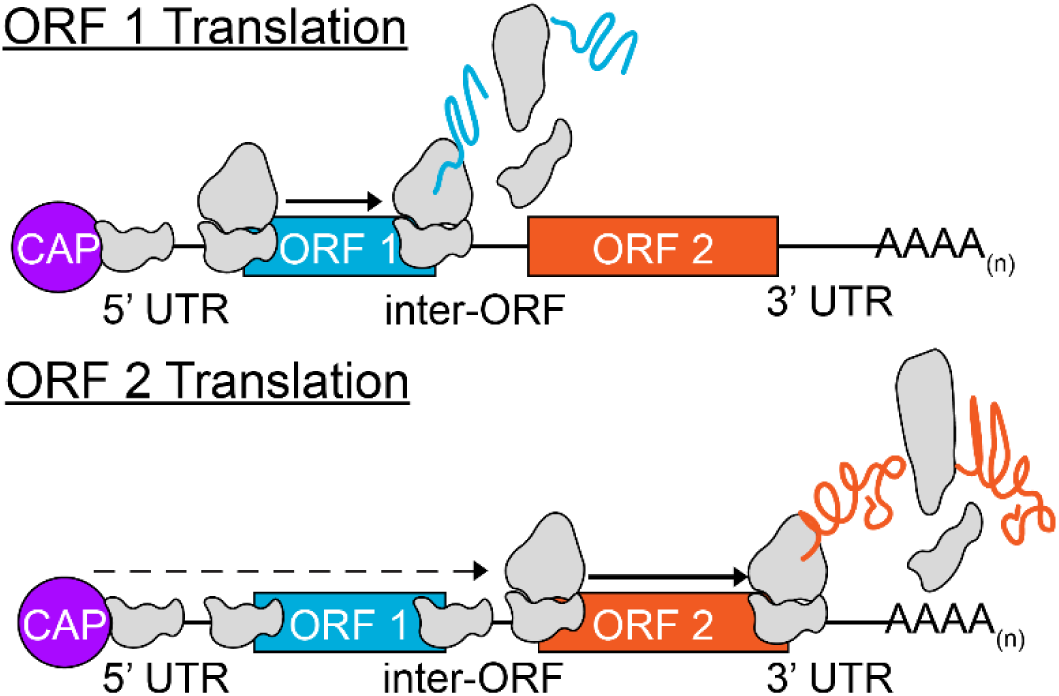
The mechanism driving bicistronic expression in green algae. Translation initiates at ORF 1 via cap-dependent scanning. Leaky scanning of ORF 1 dependent on the strength of its Kozak-like sequence allows continued scanning for subsequent translation initiation of ORF 2.

Previous efforts at bicistronic transgene expression in green algae met with limited success, and it has been difficult to test hypotheses as to the underlying mechanism. Onishi and Pringle (51) found that recovery of *Chla. reinhardtii* transformants with high expression of fluorescent reporters and fusion proteins was improved using bicistronic constructs containing the reporter gene CDS upstream, and the *aphVIII* selection marker downstream, separated by a six to 10 bp spacer sequence (51). While multiple different reporters and fusions were expressed successfully at the ORF 1 position, *aphVIII* was the only selectable marker that could be expressed at ORF 2 and yield Venus positive transformants. The authors postulated that ribosomal reinitiation, in which the ribosome complex reassembles and reinitiates translation at a downstream site after encountering a stop codon, was the most likely mechanism, but acknowledged inconsistencies with this model, specifically a documented negative correlation between uORF length and reinitiation efficiency, with the best efficiencies seen in short uORFs only a few codons in length (52, 53). This made it unlikely that reinitiation would occur after translation of the “large” ORF 1 in these constructs and did not adequately explain why the ORF 2 selectable marker gene was restricted to *aphVIII*.

Recently, Jacobebbinghaus et al. (54) integrated a series of transgenic constructs expressing bicistronic mRNAs in *Chla. reinhardtii*, and they observed that the different inter-ORF sequences alone were sufficient to produce dramatic differences in expression of both ORF 1 and ORF 2, as judged by fluorescence activity and transformation efficiency, respectively (54). The authors suggested that specific inter-ORF sequences might produce differences in mRNA secondary structure and folding energy that could influence downstream ORF translation.

Our findings are inconsistent with these models for driving bicistronic expression, as the trends seen in the mutational analyses do not reflect the expected outcome for reinitiation driving ORF 2 translation. If ORF 2 expression is dependent on reinitiation after ORF 1 translation, the strength of the Kozak-like sequence of ORF 1 should be positively correlated with ORF 2 expression, but we observed the opposite effect (Figure 5, 6). Additionally, reincorporation of a potential alternative initiation codon into the CDS of the methionine-less SuperFolder GFP would not be expected to affect *LUC* translation; instead restoring one of the AUG codons reduced luciferase activity to background levels (Figure 6, S08). The rate of reinitiation is also influenced by the spacing between ORFs. Higher rates of reinitiation are associated with inter-ORF regions of only a few bp (55), but many of the algal inter-ORF sequences are in excess of 100 bp (supplementary dataset S01), making it unlikely that substantial reinitiation could occur in these instances.

In light of these findings, the observations described by Onishi and Pringle (51) and Jacobebbinghaus et al (54) are consistent with some contribution from leaky ribosome scanning to bicistronic gene expression in their systems. The bicistronic transgene constructs analyzed in Jacobebbinghaus et al. contained multiple ATG 3-mers within ORF 1 reporter and fusion genes, features which we expect would inhibit ORF 2 translation. If only a small amount of *aphVIII* is necessary to confer resistance to aminoglycoside antibiotics this may explain why *aphVIII* was the only viable ORF 2 selection marker. We observed that, in general, the constructs analyzed by Jacobebbinghaus et al that incorporated inter-ORF sequences with fewer ATG 3-mers in the inter-ORF region had higher mVenus fluorescence and relative transformation efficiency than those with a greater number of ATG 3-mers (Figure S9). Mechanistic interpretations of these results are confounded by position effects caused by random integration of the test constructs, and the use of multiple different inter-ORF sequences, each of which changes the Kozak-like sequence governing translation of the selectable marker gene at ORF 2. In our in vivo experiments, we controlled for these variables by uncoupling reporter and selectable marker gene expression, targeting all constructs to the same (homozygous) neutral landing pad to reduce position effects, and minimizing alternative initiation codons upstream of ORF 2. Consequently, bicistronic reporter gene transcript abundances were very consistent between strains (Figure S5, S7), allowing us to conclude that changes in reporter activity were due solely to differences in translation.

Another consideration is the possibility that bicistronic transcript inter-ORF regions contain “IRES-like” elements that directly recruit ribosomes for translation of the downstream ORF. In our previous analysis, prediction software did not uncover any evidence of known IRES within polycistronic loci (27), and experimentally tested IRES from other organisms have historically performed poorly in green algae (51, 56). Onishi and Pringle (51) and Jacobebbinghaus et al. (54) both reported improved transformation efficiency (presumably via increased *aphVIII* expression) from truncated versions of inter-ORF sequences that were far too small to contain complex IRES structural elements. Our own observations also argue against IRES-like features in green algae bicistronic genes. Many algal inter-ORF regions are less than 100 nt long, with no evidence of conserved sequence motifs or structures, and some *A. protothecoides* loci even contain overlapping ORF 1 stop and ORF 2 start codons (Supplementary dataset S01). If inter-ORFs had IRES-like properties, ORF 2 translation would be expected to be independent of cap-dependent translation initiation, and modifications to the ORF 1 Kozak-like sequence should not affect ORF 2 translation. However, our in vivo mutational analyses of endogenous and synthetic bicistronic loci demonstrated that ORF 2 translation was directly influenced by sequence manipulations at the ORF 1 initiation site (Figures 5 and 6), contradicting the IRES hypothesis. We conclude that the inter-ORF sequences do not contain IRES-like elements, but rather 5’ UTR length, Kozak-like strength, and the frequency of alternative translation initiation sites upstream of ORF 2 determine the amount of leaky scanning and ability of the ribosome to reach ORF 2, which may explain the limited success in achieving bicistronic gene expression in previous studies (51, 54).

### Regulation of bicistronic gene expression

The ability of the 43S pre-initiation complex to recognize a start codon within the proper sequence context is largely conferred by eukaryotic initiation factor 1 (eIF1) (57). eIF1 promotes the transition from an “open”, scanning-competent preinitiation complex to a “closed” state that interacts with the Kozak-like sequence and initiates translation (58). The translational landscape is profoundly altered in *eIF1* knockdown and knockout human cell lines, with ribosome footprints that reveal far less discrimination between canonical translation start sites and alternative upstream initiation sites with poor Kozak sequences (59). Intracellular eIF1 concentrations can be influenced by stress response conditions, suggesting that this may provide a mechanism for regulating ORF expression ratios from algal bicistronic transcripts (59, 60). Intriguingly, genes with altered translation in *eIF1* knockdown lines were enriched for functions in energy production and sensing of metabolic stress; including targets such as TOM subunits and proteins with LYR- motifs, both of which are seen in conserved bicistronic genes in green algae (27). In addition to a subpar Kozak-like sequence around the start codon, hairpin structures around the initiation site have been shown to influence the recognition efficiency of the 43S pre-initiation complex and may promote leaky scanning in specific scenarios where the ORF 1 Kozak-like context is strong (56).

Another level of bicistronic gene regulation is provided by nonsense mediated decay (NMD), a quality control mechanism that typically degrades aberrant mRNAs containing premature termination codons (61). In *A. thaliana*, mRNA transcripts encoding uORFs longer than 50 amino acids (within the size range of ORF 1s in algal bicistronic transcripts) are targets of the NMD pathway (62). The *CDC26/TTM3* locus, encoding a 65 amino acid cell cycle regulator at ORF 1 and an inorganic polyphosphatase at ORF 2, is conserved in both plants and green algae, and illustrates the interplay between NMD and translation (11). *CDC26/TTM3* transcript and gene product levels were higher in *A. thaliana* NMD mutants compared to wild-type, suggesting that abundance of the bicistronic mRNA is regulated by NMD, triggered by the short *CDC26* ORF (11). Conversely, translation of the *TTM3* ORF enhanced *CDC26* ORF 1 protein levels by recruiting the bicistronic transcript to polysomes, suggesting a mechanism for bypassing NMD to improve production of small proteins (11). It is possible that this may be a more general regulatory mechanism that extends to green algal bicistronic genes.

### Why leaky ribosome scanning for bicistronic expression?

Polycistronic gene arrangements increase physical compactness of the genome and guarantee coordinated gene expression. Leaky ribosome scanning is a commonly used mechanism for translating viral polycistronic transcripts (19). For example, the plant pathogen *Plantago asiatica* mosaic virus (PlAMV) utilizes leaky scanning to translate three “triple gene block” proteins (TGBp) involved in viral genome transport from a tricistronic mRNA (34). The Human Papillomavirus (HPV) E6/E7 bicistronic mRNA displays an extreme degree of leaky scanning, requiring bypass of 12 internal AUG 3-mers in the E6 ORF in order to translate the E7 ORF (23). Given their genome size constraints, it is understandable that viruses employ leaky scanning for coordinated protein synthesis, but even the smallest green algal genomes are orders of magnitude larger than those of viruses. Alternatively, leaky scanning can contribute to dual functionality of a transcript via production of alternative protein isoforms. A suboptimal Kozak-like context sequence around the first AUG of the *S. cerevisiae PiF1* transcript, encoding a helicase, permits leaky scanning, resulting in translation initiation at a downstream AUG to produce a truncated protein isoform that is targeted to the nucleus instead of the mitochondrion (63). In *A. thaliana*, leaky scanning of the DNA polymerase-encoding *POLγ2* mRNA similarly generates alternative protein isoforms with targeting to either the chloroplast or the mitochondrion (64). Although some algal bicistronic transcripts have splice variants (27), we have not encountered examples encoding protein variants with alternative organellar targeting signals. Instead, some bicistronic genes may coexpress proteins that function together. However, this does not appear to hold for a majority of bicistronic genes, as functional predictions suggest that many encode pairs of proteins with different localizations and unrelated functions (Supplementary dataset S04). We note that about a quarter to a third of proteins encoded at bicistronic loci are predicted to be localized in either the chloroplast or mitochondrion (Figure S10 and supplementary dataset S04). This is particularly intriguing in that polycistronic expression is often found in organellular genomes, suggesting that these nuclear genes may have ancestral plastid or mitochondrial origin.

uORFs are ubiquitous throughout eukaryotic genomes (65). Nearly 67% of annotated *Chla. reinhardtii* transcripts contain at least one uORF (38); they are present in more than half of the annotated transcripts in humans and mice (66), 30-70% of plant mRNAs (46, 67), and thousands of candidate uORFs are present in *S. cerevisiae* genes (68). Their primary role appears to be inhibition of 5’ cap-dependent translation of canonical protein coding ORFs (40, 66), where translation of canonical ORFs can proceed through alternative translation initiation events such as ribosome reinitiation and leaky scanning. Whether these uORFs encode functional polypeptides remains a question, but it is evident that bicistronic transcripts in green algae occur as a continuum with the phenomenon of uORFs. The first ORF in green algal bicistronic genes can be considered to be a “large” protein coding uORF, and like other uORFs, appear to inhibit the translation of downstream ORFs (Figure 4, 5, and 6). But outside of this regulatory role, both ORFs of most algal bicistronic loci are highly conserved and/or have predicted functionality. This further supports our premise that polypeptide-encoding uORFs may be more prevalent in eukaryotes than previously thought.

### The functional role of proteins from bicistronic loci

To better determine the biological significance of proteins encoded by bicistronic transcripts, functional and localization predictions were performed for both previously identified and novel genes. Over one-fourth and one-seventh of proteins encoded by bicistronic loci in *A. protothecoides* and *Chla. reinhardtii* respectively, are predicted to have localization or function associated with the mitochondria (Figure S10 & Supplementary Dataset S05). Cre06.g278242/Cre06.g278245 and UTEX250_A10.30135/ UTEX250_A10.30130) encode TOM22, an ortholog of Arabidopsis TOM9 at ORF 1 predicted to aid in translocase of the outer membrane (TOM) complex assembly, and SDHAF3 at ORF 2 with predicted function in succinate dehydrogenase assembly for respiration. This locus is widely conserved in bicistronic fashion throughout green algae (27). The UTEX250_A05.08765/ UTEX250_A05.08770 locus encodes proteins with predicted functions in mitochondrial respiratory chain assembly and a cytochrome *c* oxidase assembly factor. In fact, 5/22 of the *A. protothecoides* bicistronic loci with mitochondrial association encode proteins with functions in cytochrome *c* oxidase assembly. UTEX250_A12.36865 is an ortholog of human PET100, functions in the biogenesis of Complex IV and is a component of the mitochondrial electron transport chain (69). About 20% of the genes in *A. protothecoides* and 10% in *Chla. reinhardtii* are predicted to encode proteins that function and localize to the nucleus. Notably, the *REX1S-REX1B* locus, which has been experimentally analyzed in *Chla. reinhardtii*, produces a bicistronic mRNA encoding two polypeptide subunits thought to function in DNA repair (70). Also found in bicistronic context are proteins predicted to function in the secretory pathway. UTEX250_A05.11130/UTEX250_A05.11130 & Cre16.g801965/Cre16.g686900 encode OST4, a 33 amino acid subunit of an oligosaccharyltransferase complex for ER polypeptide modification, which have homologs in yeast (71). A sizable number of proteins are predicted to have localization, or functional domains associated with the chloroplast. UTEX250_A10.29670, conserved in *Chla. reinhardtii*, is an ortholog of cyclophilin 37, associated with maintaining the stability of electron transport chain in photosynthesis under high light stress (72). Despite evidence of their conservation and translation, many bicistronic genes encode proteins of unknown function, with 29 predicted ORFs in *A. protothecoides* and 16 in *Chla. reinhardtii* having no known orthologs or functional predictions (Figure S10 & Supplementary Dataset S05). These “pioneer” proteins present candidates for future reverse genetic analysis for novel gene discovery.

### Application and utility in synthetic biology

Green algae are promising microbial systems for synthetic biology and environmental applications such as bioremediation (26). Engineering in these systems requires expression of multiple transgenes to build synthetic biochemical pathways, and polycistronic arrangements may be useful for coordinating gene expression and modulating protein stoichiometry. Leaky ribosome scanning has been successfully used in other organisms for transgene expression. Tunable coexpression of three ORFs and successful production of monoclonal antibody components was achieved in transient assays in human cell lines by modifying the order and the strength of the Kozak-like sequence for each ORF (48). Although leaky scanning imposes several constraints, it may prove to be an advantageous strategy for production of proteins with specific stoichiometries. Additionally, the recruitment of polysomes at bicistronic loci (11) could be of particular interest for improving the translation of small polypeptides that might otherwise be difficult to produce. Leaky scanning may be widely applicable because the process of translation initiation is itself widely conserved in eukaryotes. That said, using this strategy should employ multiple design considerations, including a short 5’ UTR, removal or limiting AUG sequences upstream of ORF 2, a sub-optimal Kozak-like sequence at ORF 1, and a strong Kozak-like sequence at ORF 2. Outside of the context of polycistronic expression, manipulation of the Kozak-like sequence is a powerful tool for adjusting protein output without changing transcript abundance. In rabbits, manipulation of the Kozak-like sequence context of the *PCSK9* gene involved in cholesterol homeostasis induced phenotypic variations without influencing transcription (73). These insights can guide construct design for transgene expression and encourage studies to further optimize expression of multiple ORFs for use in synthetic biology.

## Materials and Methods

### Manual curation of bicistronic genes

The transcriptomes and genomes of *Auxenochlorella protothecoides* (UTEX250-A *v1.0*) and *Chlamydomonas reinhardtii* (*CC-4532 v6.1*) were visualized on the Integrative Genomics Viewer (74) with input from Iso-Seq and RNA-Seq data. For *Chla. reinhardtii*, additional datasets for H3K4me3 ChIP-Seq, Ribo-Seq, and DNA consensus sequences for field strains CC-1952 and CC2931 were included (29). Bicistronic loci for both algae were characterized in terms of 5’ UTR size, ORF size, ORF spacing, coexpression relative to colinear genes, and conservation in other Chlorophyte species. A detailed description of the high-confidence evaluation and gene structure properties can be found in supplementary dataset 1 (S01).

### Bicistronic loci Conservation Analysis

Criteria for conservation was adapted from our previous work with minor modifications (31). The protein sequences of all ORFs in the bicistronic loci were used as query searches for protein-protein similarity search in other green algae, whose genomes were available from either Phytozome.net (*Chlamydomonas reinhardtii* v6.1, *Chromochloris zofingiensis v5.2.3.2, Coccomyxa subellipsoidea* C-169 v2.0, *Dunaliella salina* v1.0, *Ostreococcus lucimarinus* v2.0, *Micromonas pusilla* CCMP1545 v3.0, and *Volvox carteri* v2.1) or Phycocosm.net (*Chlorella Sorokiniana* UTEX 1602, *Chlorella vulgaris*, and *Nannochloropsis desiccata* UTEX 2437) True hits were determined using a BLAST bit score cutoff of ≥30. This database was then queried to identify pairs of colinear ORFs (ORFs on the same reading strand separated by <10 kbp) as possible bicistronic loci. The list of candidates was then manually curated using either the JBrowse tool from Phytozome.net or visualization in Geneious Prime. When available, expressed sequence tag (EST) data from Phytozome was used to identify transcripts spanning two or more ORFs in candidate polycistronic loci. Similarly, Iso-Seq data from *Chlamydomonas reinhardtii*, *Chromochloris zofingiensis*, *Dunaliela salina*, and *Chlorella sorokiniana* were used to determine if colinear ORFs are transcribed from bicistronic mRNA in those species. Additionally, some bicistronic orthologs in *Chla. Reinhardtii* were previously identified using the OrthoFinder software (75). For these hits, orthologs were manually viewed on IGV to ensure they were authentic bicistrons. For each pair of hits in each species with a high scoring pair, the protein ID (or "unannotated" for putative proteins encoded by unannotated ORFs), the coordinates of the gene encoding that protein in the corresponding genome assembly, the BIT score of the match, supporting evidence (either colinear ORFs, EST support, or Iso-Seq support) and the BLAST algorithm used can be found in Supplementary dataset 2 (S02).

### Bicistronic Gene Protein Prediction

The protein sequences of all ORFs in the bicistronic loci were exported as a FASTA file and analyzed for Pfam domains and GO Ontology using InterPro 100.0 (https://www.ebi.ac.uk/interpro/). Subcellular localization predictions for all proteins were determined using the DeepLoc 2.0 software (https://services.healthtech.dtu.dk/services/DeepLoc-2.0) Associated accessions, Pfam domain matches, InterPro and PANTHER GO terms, DeepLoc prediction scores, and any additional notes can be found in supplementary dataset 4 (S04).

### Kozak-like Sequence Generation and Kozak Bit Score Comparison

Kozak scores for monocistronic and bicistronic genes were quantified using previously described methods (38,39). For *Chla. reinhardtii*, monocistronic nuclear gene models from the v6.1 annotation were first filtered to remove transposable element genes and any genes that did have at least one homolog in a representative collection of high quality Chlorophyceae and Trebouxiophyceae genomes (Craig et al, unpublished). Next, the filtered monocistronic gene set (N=14,738) was randomly divided into two datasets containing an equal number of genes. The first random division was used to generate a consensus Kozak logo using WebLogo 3 (39) by extracting the 5 bp upstream and downstream of each annotated start codon. A Kozak score was then calculated for each of the genes in the second half of the monocistronic dataset, and for all the bicistronic genes. In the consensus logo, each of the four nucleotides at each site in the Kozak sequence has an associated bit score. The Kozak score is the sum of bit scores for each nucleotide in a query sequence based on the corresponding nucleotide in the consensus logo. The same method was applied to *A. protothecoides* strain UTEX 250-A. Nuclear monocistronic genes were filtered to remove genes without homologs, and the haplotype A gene model was arbitrarily selected when a gene was represented by two alleles in the diploid genome. The filtered dataset (N=7,648) was divided randomly in two, a Kozak consensus logo was generated from one half, and Kozak scores were generated from the other half and for the bicistronic genes.

### ATG 3-mer analysis

To estimate the bias of “ATG” codon frequencies within a given sequence, it was established that given a sequence of length *n*, there is a 0.25 probability for each nucleotide to be encountered at any individual position in a sequence with 50% GC content. Hence, the probability of encountering an “ATG” sequence by random chance is 0.015625 (0.25 *0.25 *0.25). The probabilities for G and C were further adjusted by dividing 0.25 by the respective GC content (64.1% for *Chla. reinhardtii* and 63.8% for *A. protothecoides*) to better account for codon bias. The adjusted probabilities (*p*) for encountering an “ATG” in any random 3-mer were then calculated to be 0.01032659 (0.1795 * 0.1795 * 0.3205) and 0.010450759 (0.181 * 0.181 * 0.319) for *Chla. reinhardtii* and *A. protothecoides* respectively. FASTA files were then generated containing sequences for the following specified regions of bicistronic loci: 5’ UTR, ORF 1, inter-ORF, and ORF 2. These were also generated for the ORFs of all monocistronic genes with an annotated 5’ UTR (n = 12,232 in *Chla. reinhardtii*, n =5180 in *A. protothecoides*) in addition to a separate FASTA containing only the sequences of the 5’ UTRs. Using an R script, individual sequence lengths and total “ATG” 3-mers were calculated for every sequence of the dataset. To account for a start codon where translation would canonically occur, a value of 1 was subtracted from any sequence annotated as an ORF. Calculated lengths and codon counts were further authenticated via manual curation of all bicistronic genes and 50 random genes from the monocistronic dataset. Using the formula [(*n* – 2) * *p*], the expected “ATG” frequency (*e*) was calculated for every sequence in the dataset. The actual observed value (*o*) obtained from the R script was then divided by the estimation to create a ratio representing the “ATG” bias (*o*/*e*) of a sequence based on its length. A detailed dataset containing the calculation template, sequence lengths, observed and estimated “ATG” counts, and the observed/expected values for all sequences used in this study can be found in supplementary dataset 3 (S03).

### PCR, Cloning, Plasmid Assembly, Cassette Construction

A detailed description of all primers, templates, vectors, plasmids, and assembly instructions used in this study can be found in supplementary dataset 5 (S05). In general, primers for fragment amplification were designed using the NEBuilder® Assembly Tool (https://nebuilder.neb.com/) and ordered from Integrated DNA Technologies (IDT, San Diego, CA). Codon optimization was performed according to a codon usage table from *Prototheca moriformis* UTEX 1435 (US Patent US7935515B2) using *Gene Designer* (ATUM Bio Inc., Newark, NJ) software. Codon usage was based on the most frequently used codon, substituting less frequently used codons to avoid unwanted restriction sites, repeats and cryptic splice donor or acceptor sites. The *Venus* CDS was amplified from plasmid pLM005 (76) and then incorporated into other templates as described in S04. The codon-optimized Gaussia princeps LUC CDS was synthesized by GenScript Biotech, (Piscataway, NJ) All in silico cloning work was conducted using Geneious Prime by Dotmatics (Boston, MA).

PCR was performed using Herculase II Fusion DNA Polymerase (Agilent #600679). Thermocycler conditions were set according to the manufacturer’s recommendations based on concentrations, target sequence length, and primer Tm (available in S04). PCR products were purified with an Omega Bio-Tek E.Z.N.A.® Gel Extraction Kit. Fragments and linearized vector backbones were assembled using the NEBuilder® HiFi DNA Assembly mix and introduced into E. cloni® 10G Chemically Competent Cells using a 30 second heat shock method. Colonies were picked and grown overnight at 37°C with 250 rpm shaking in 2 mL of Luria Broth (LB) media with a concentration of 200 μg/mL ampicillin for selection. Plasmid purifications were conducted with an E.Z.N.A.® DNA Plasmid Mini Kit I (Omega Bio-Tek Inc. Norcross, GA), and concentrations were determined via NanoDrop (Thermo-Scientific NanoDrop^TM^ One). Correct assembly and sequence were validated through Nanopore and Sanger sequencing (UC Berkeley DNA Sequencing Facility). For scale-up, 1-2 uL of verified plasmid minipreps were transformed into 10G competent cells and cultures were grown overnight at 37°C with 200-250 rpm shaking in 200 mL LB with 200 μg/mL ampicillin. Plasmid DNA maxipreps were performed using an E.Z.N.A.® DNA FastFilter Maxi Kit (Omega Bio-Tek Inc. Norcross, GA) per the manufacturer’s instructions. DNA concentrations were measured with a NanoDrop One spectrophotometer (Thermo Fisher Scientific, Waltham, MA).

### *A. protothecoides* strain, media, and culture conditions

*A. protothecoides* strain UTEX250 was obtained from the UTEX Culture Collection of Algae (The University of Texas at Austin). ApM1 culture media recipe was adapted from a composition described in US Patents US20140178950A1 (pg. 28) and US Patent US8927522B2 (pg. 37). Per liter of MQ-grade water, the following amounts of salts were added for the following concentrations: 4.2 g (24.1 mM) potassium phosphate dibasic anhydrous (Fisher P288-500), 3.57 g (25.9 mM) sodium phosphate monobasic monohydrate (Fisher S369-1), 240mg (974 µM) magnesium sulfate heptahydrate (Fisher M63- 500), 250 mg (1.3 mM) citric acid (Fisher BP339-500), 1.7 mL (170 μM) of a 100 mM calcium chloride dihydrate stock solution (Fisher C79-500), 2 μM thiamine-HCl (Sigma T1270-25G), and 10 mL of 100X Ap trace element solution. A 1 L stock of 100X Ap trace element solution consisted of 2.743 g (14.3 mM) citric acid (Fisher BP339-500), 11 mg (14.3 μM) copper sulfate pentahydrate (Fisher BP346-500), 330 mg (340.5 μM) boric acid (Sigma B9645-500G), 1.4 g (4.89 mM) zinc sulfate heptahydrate (Sigma Z0251-500G), 948.5 mg (4.79 mM) manganese chloride tetrahydrate (Sigma M3634-100G), 3.9 mg (161 μM) sodium molybdate dihydrate (Sigma M1651-100G), and 110 mg (396 μM) iron sulfate heptahydrate (MP Biochemicals 194663). For solid media used in colony selection, 1.5% (w/v) agar (Fisher BP1423-500) was added to MilliQ water and then autoclaved before mixing with the other solutions to achieve correct concentrations. Cell cultures were started from a small loop or colony of the strain of interest and grown in either photoautotrophic or heterotrophic conditions. For photoautotrophic conditions, cultures were grown in ApM1 liquid media with the addition of .5% (w/v) glucose and 2 μM thiamine-HCl. Cultures were grown at 26C and 50–100 µmol photons/m^2^/s and shaken continuously at 140 revolutions per minute (RPM). For heterotrophic conditions, cultures were grown with ApM1 media with additional 2% (w/v) glucose. These cultures were also shaken continuously at 140 revolutions per minute (RPM) but were grown in complete darkness. For transformant selection in either liquid culture or agar plates, G418 Sulfate (VWR E859-5G) was added at a concentration of 100 µg/mL.

### *A. protothecoides* transformation and phenotyping

Lithium acetate transformation was adapted from the protocol described in US Patent US- 12037630-B2. 1M stock solutions of lithium acetate (Acros Organics, #6109-17-4), tris-HCl (Fisher BP153- 500), and polyethylene glycol 4,000 (PEG-4000, Alfa Aesar A16151) were made before starting the transformation. For cassette targeting, 100 μg of plasmid DNA diluted to a 750 μL solution was linearized by restriction digest with XbaI, (New England Biolabs) using recognition sites flanking the *DAO1* 5’ and 3’ targeting sequences. Digests were then extracted with 750 μL of 25:24:1 phenol-chloroform-isoamyl alcohol pH 8 (Fisher #BP17521-400) and centrifuged at maximum speed for 10 minutes in a tabletop centrifuge (Eppendorf). The aqueous phase was transferred to a new 1.5 mL eppendorf tubes, mixed with 525 μL isopropanol (Sigma-Aldrich W292907), and incubated at room temperature for 1 hour to precipitate the digested DNA. DNA was pelleted by centrifugation at maximum speed for 30 minutes; pellets were washed twice with 500 μL of 70% Ethanol (Sigma-Aldrich E7023) spinning for 5 minutes at maximum speed between each wash, and then dried in a sterile laminar flow hood. Linearized plasmid DNA was dissolved in 50-100 μL of elution buffer (10 mM Tris-HCl, pH 7.5) (Omega Bio-Tek Inc. Norcross, GA).

To prepare cells for transformation, a loop of UTEX 250 wild-type cells from a plate culture was inoculated in a 50 mL ApM1 media with 2% (w/v) glucose. Cultures were grown at 28°C in complete darkness with shaking at 140 rpm until an OD750 between 1.75-3.5 was reached. Cells were then transferred to 50 Polypropylene Conical Tube (Falcon 352098) and then pelleted by centrifugation at 3750 rpm for 5 minutes. Cells were then washed in 5 mL of a 0.1M lithium acetate/1X Tris EDTA (TE) solution and centrifuged at max speed. The cells were then resuspended in 500 μL 0.1M lithium acetate/1X Tris EDTA (TE) solution and allowed to sit shake at 200 rpm for one hour. Cells were then separated into 150 uL aliquots and incubated with 15 μg of linearized DNA for 30 minutes. Next, 750 μL of 0.1M lithium acetate/1X Tris EDTA (TE) / 40% PEG-4000 was added for an overnight incubation at room temperature. The next day, cells were collected via centrifugation and placed in ApM1 with 2% glucose and 1X thiamine media for an 8-hour recovery period. Afterward, cells were centrifuged to form a pellet, resuspended in 250 μL of 1M sorbitol, and plated onto ApM1 (with 2% (w/v) glucose, 2 μM thiamine-HCl, and 1.5% agar) selection plates with a concentration of 100 ng/μL G418. After a 7-to-14-day dark incubation period at 26C, colonies were picked and grown heterotrophically in 1 mL ApM1 liquid cultures with 2% (w/v) glucose, 2 μM thiamine-HCl, and 100 ug/mL of G418 for selection. After 3-5 days, a 1/2500 serial dilution of the culture was conducted before replating on selective media for single colony isolation. Colonies that grew on these plates were further isolated for further analysis.

For genotyping, DNA from wild type UTEX 250 cells and transformant strains was extracted using a CTAB extraction method adapted from European Patent EP2785835A2. 2mL of wild type UTEX 250 cells were grown to stationary phase and then placed in a screw-capped tube, centrifuged at maximum speed, and then frozen at -20C with a 4mm glass bead (Fisher 11-312B). A mixture of 300uL grinding buffer (adapted from European Patent EP2785835A2), 1.5 μL RNase A (VWR E866-1ML), and 250 mg of 0.5 mm glass beads (Fisher35-535) was added, and the tube was shaken for 5 minutes in a bead beater. To this, 40 μL of 5M NaCl (VWR E529-500ML) and 66 μL 5% CTAB (VWR 0833-500G) were added, mixed by inversion, and incubated at 65C for 1 hour. After incubation, samples were centrifuged at maximum speed for 10 minutes, and the supernatant was transferred to a fresh tube. To this, 300 μL of 25:24:1 phenol-chloroform-isoamyl alcohol pH 8 (Fisher BP17521-400) was added before centrifugation at maximum speed for 10 minutes. The aqueous phase was transferred to a new tube and mixed with 210 μL of isopropanol and incubated at room temperature for 1 hour. This mix was then centrifuged at max speed for 30 minutes to form a pellet. The pellet was then washed with 2 rounds of 500 μL 70% EtOH and dissolved in elution buffer (10 mM Tris-HCl. pH 7.5). DNA nanodrop concentrations were further diluted to have ideal genomic DNA concentrations around 100-300 ng/μL for use with the Herculase II Fusion DNA Polymerase (Agilent 600679). A detailed description of genotyping primers can be found in supplementary dataset 5 (S05).

### RNA extraction and RT-qPCR analyses

For the collection of RNA, 50 mL photoautotrophic and heterotrophic cell cultures were harvested during mid-log phase of growth (∼2.5 days). RNA extraction was carried out using the ZymoBIOMICS^TM^ RNA Mini Kit (Zymo, R2001) per the manufacturer’s protocol. To generate cDNA, 2.5 μg of extracted RNA were mixed with 1 μL of 10 mM dNTP mix (New England Biolab), and 1 μL of 500 μg/mL oligo (dT)18 (New England Biolab), then incubated at 65°C for 5 minutes. Reactions were cooled on ice, and a mix containing 4 μL first-strand buffer, 2 μL of 0.1 DTT (New England Biolab), 1 μL of RNasin (New England Biolab), and 1 μL M-MLV (New England Biolab) were added to each sample. This was followed by a 50-minute incubation at 42°C and a 15-minute heat shock at 70°C. The resulting cDNA was then diluted 10-fold with MilliQ-grade H2O.

RT-qPCR primers specific to reporter CDS sequences were designed with the PrimerQuest™ Tool from Integrated Gene Technologies (IDT) emphasizing amplicon products less than 200 bp and proximity toward the 3’ end of the CDS. RT-qPCR reactions were conducted in 96-well plates using a CFX96 Real-Time System (BIO-RAD, Berkeley CA). A 20 μL reaction mix, consisting of 4 μL cDNA, 10 μL iTaq Universal SYBR® Green Supermix (BIO-RAD, Berkeley CA), and 2 μL each of reverse and forward primers, was added per well. 2-step amplification and melting curve cycles were repeated 39 times with denaturing step at 95°C and annealing/extension at 60°C. Primer efficiencies were resolved with a template sequence and five 10-fold dilutions. For absolute abundance, a plasmid (pMAD57) of known concentration, was used as a template for seven 10-fold dilutions. The molar concentration of the template was converted to log2 molecules and plotted against the Ct values to construct a standard dilution curve. The Ct values for each mRNA sample were then inputted into the line of best fit equation to calculate transcript abundance. All samples were run alongside a no-template control and a no-reverse transcriptase control. A detailed description of all RT-PCR and RT-qPCR primers can be found in supplementary dataset 4 (S04).

### Fluorescence and luminescence quantification and visualization

Photoautotrophic and heterotrophic cells were grown in 1 mL 24 well plates for a period of 128 hours before phenotyping, at which point OD750 was measured for normalization with a Spectramax iD3 (Molecular Devices, LLC., San Jose, CA) well plate reader, which was also used for subsequent fluorescence and luminescence measurements. Before taking a measurement, cell plates were set to shake at a speed of “high” for 5 seconds. Venus fluorescence was measured using an excitation-emission spectra of 515/555. Superfolder GFP was measured using an excitation-emission spectra of 488/528. Luciferase readings were conducted using the Pierce™ Gaussia Luciferase Flash Assay Kit. Working coelenterazine solution was made per manufacturer’s instructions. For luminescence quantification, a 20uL sample of cells were taken after well plate shaking and loaded into a black, clear bottom 96 well plate. A 50uL working solution was then added to each biological replicate and relative luminescence was measured with the Spectramax iD3 at 487 nm. For visualization of luminescence, 200uL of working solution was added to a 1mL culture of cells and gently mixed for a period of 10 minutes. Visualization was conducted using an Azure 200 Gel Imaging Workstation under the “chemiluminescence” setting with manual capture set to a period of 3 minutes.

For confocal microscopy images, 50uL aliquots were taken and centrifuged at a speed of 10,000g for 1 minute. Cells were then resuspended in 50uL of a 10uM Sodium Phosphate Buffer Solution (pH =7). 10uL of each sample were then put on Fluorescent Antibody Slides (Thermo Scientific Cat. No. 3032) and covered with a Microscope Cover Glass (Fisherbrand 12-542A) and sealed with nail polish. Samples were observed under a Zeiss LSM880 Laser Scanning Confocal Microscope with a 63x oil objective lens (numerical aperture 1.15). Venus Signal was observed with a laser excitation line at 514 nm (intensity, 20%); emission was collected between 519-599 (gain 650; offset 0). Superfolder GFP Signal was observed with a laser excitation line at 488 nm (intensity, 20%); emission was collected between 510-550 (gain 700; offset 0). Chlorophyll Fluorescence was observed with a laser excitation line at 635 nm (intensity, 4%); emission was collected between 647-721 (gain 650; offset 0). Corresponding brightfield images were acquired with a transmitted detection module with a Photon Counting PMT (T-PMT, gain 371; offset 0). Images were processed and color corrected using ImageJ software. The Brightness and Contrast range for Chlorophyll was set between 10 and 150. The Brightness and Contrast range for Superfolder GFP was set between 0 and 100.

### Analysis from Jacobebbinghaus et al. 2024

In a previous study, the inter-ORF sequences taken from 22 bicistronic loci in *Chla. reinhardtii* were inserted between an *mVenus* reporter (ORF 1) and a promoter-less *aphVIII* drug-selectable gene (ORF 2) in a bicistronic expression vector (Jacobebbinghaus et al 2024). These expression vectors were then used to transform *Chla. reinhardtii*, placed under drug selection, and scored for *mVenus* expression to evaluate which if these candidate sequences (CSs) could facilitate the co-expression of both *mVenus* and *aphVIII* bicistronically. In this work, we reanalyzed that data to determine what contribution ORFs internal to the 22 candidate sequences (referred to in that work as CS1 - CS22) might have on co-expression. For each CS, a score was calculated as follows: First, the *mVenus* signal intensity from ORF 1 was summed from the percentage of transformants with low, medium, and high signal intensity as depicted in *Jacobebbinghaus* et al. Fig. 1b. CS22, which was reported as "n.a." was assigned a signal intensity of 0. Second, the transformation efficiency was determined from the length of each bar as depicted in Jacobebbinghaus et al. Fig. 1c. Finally, an efficiency score was calculated by multiplying these two values. The 22 candidate sequences (Jacobebbinghaus et al. SI Data S1) were evaluated for the presence of additional ORFs. The 22 CSs were then sorted into six categories as follows: 1) those with no additional ORFs in the CS (n = 6), 2) those with an ORF that begins and ends within the CS (n = 2), 3) those with an ORF that is in-frame with ORF 2 (n = 4), 4) those with an ORF that is 1 nt out of frame relative to ORF 2 (n = 1), 5) those with an ORF that is 2 nt out of frame relative to ORF 2 (n = 4), and 6) those with multiple and overlapping ORFs (n = 5).

## Acknowledgments

This work was supported by the Department of Energy (DOE) Office of Science, Biological and Environmental Research program under award no. DE-SC0023027 (to SSM and JLM). M.A.D was supported, in part, by the National Institutes of Health (NIH) T32 Genetics Dissection of Cells and Organisms Grant (1T32GM132022-01) and the Newton Graduate Fellowship in Synthetic Biology (QB3- Berkeley). Confocal fluorescence microscopy was performed at the CRL Molecular Imaging Center, RRID:SCR_017852. We thank Holly Aaron, Feather Ives, and Luis Alvarez for their microscopy advice and support. We thank Kyle J. Lauersen for providing a template plasmid for the *Gaussia princeps* Luciferase. We also extend thanks to Vincent Leon Gotsmann and Felix Willmund for their *Chla. reinhardtii* Ribo-Seq dataset.

## Author Contributions

M.A.D, S.D.G, R.J.C, J.L.M, and S.S.M. designed research; M.A.D and J.L.M performed cloning, cell culturing, transformation, genotyping, RT-qPCR analysis, and data collection. M.A.D, S.D.G., and R.J.C performed bioinformatic analysis; M.A.D, S.D.G, R.J.C, J.LM, and S.S.M. prepared and edited the article.

