## Supplementary Information for "Leaky ribosomal scanning enables tunable translation of bicistronic ORFs in green algae"

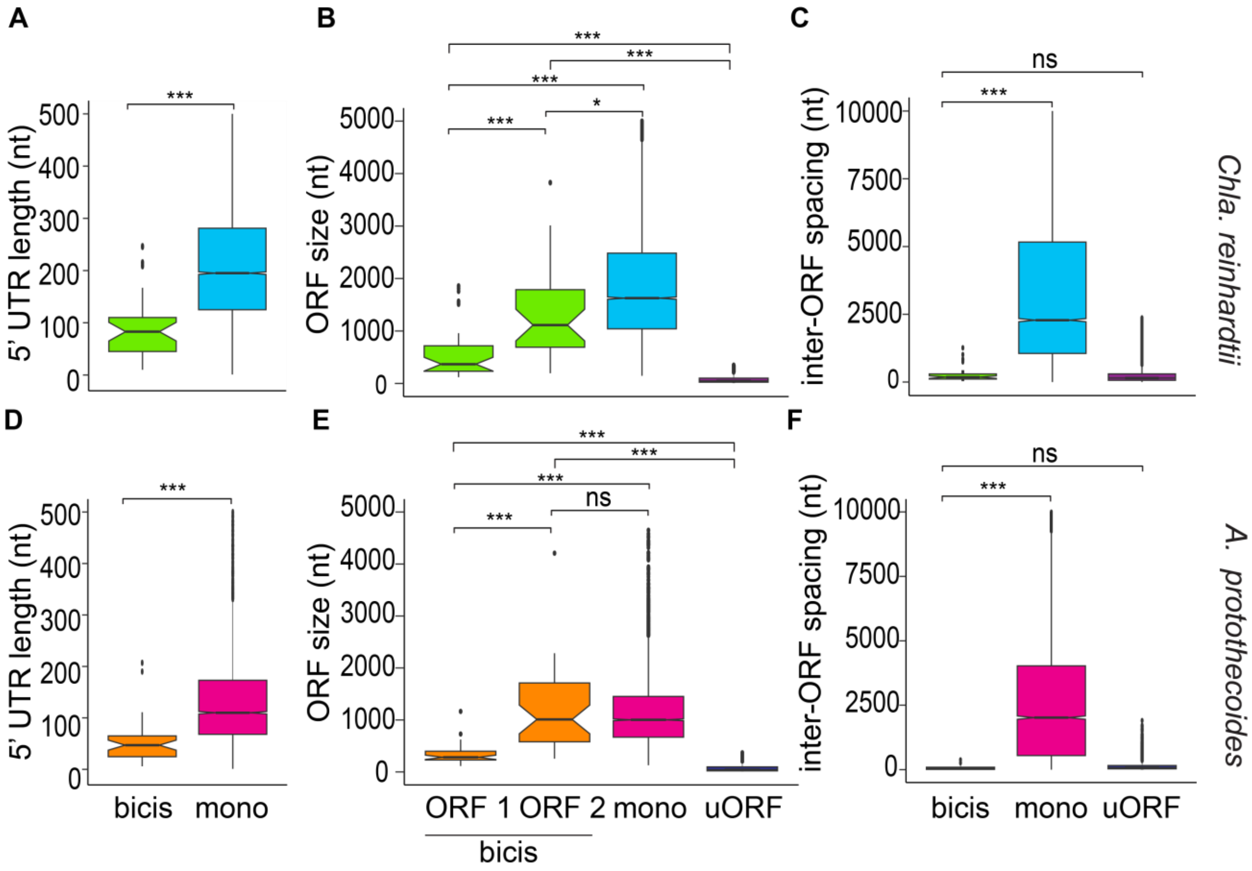


**Figure S1. Structural features of bicistronic loci.**

**(A and D)** The distribution of 5' UTR length in nucleotides. Highly conserved bicistronic genes (bicis) and protein coding monocistronic genes (mono). In *Chla. reinhardtii*, a sample size of n = 35 for bicistronic genes and n = 15,333 genes was used. In *A. protothecoides*, a sample size of n = 41 and n = 7698 was used. (**B and E)** Sizes of the protein coding ORFs in bicistronic genes (n =35, 41), monocistronic genes (n = 15,333, 7698), and uORFs ( n= 31774, 54525) in *Chla. reinhardtii* and *A. protothecoides* respectively. Bicistronic genes are separated based on their orientation as the upstream (ORF 1) or downstream (ORF 2) ORF relative to the 5' end of the transcript. (**C and F)** Distance between the stop and start codons of colinear gene pairs denoted as the "inter-ORF spacing" in *Chla. reinhardtii* and *A. protothecoides* respectively. The first plot denotes the length for all highly conserved bicistronic genes (n=35, 41) genes. The second denotes colinear (adjacent genes on the same strand of the same chromosome with ≤ 20,000 nt between ORFs) monocistronic genes (mono, n =12,213, 7698), and the third denotes uORFs within the 5' UTR of the transcript (uORFs, n = 29,210, 4457). For all plots, whiskers indicate 1.5 times the interquartile range and notches indicate the confidence interval of the median. Outliers are plotted as individual points. Statistical significance was tested for with a Kruskal-Wallis test and Wilcoxon rank-sum test for sample comparison. Asterisks "*", "**", and "***" indicate *p*-values less than .05, .01, and .0001 respectively.

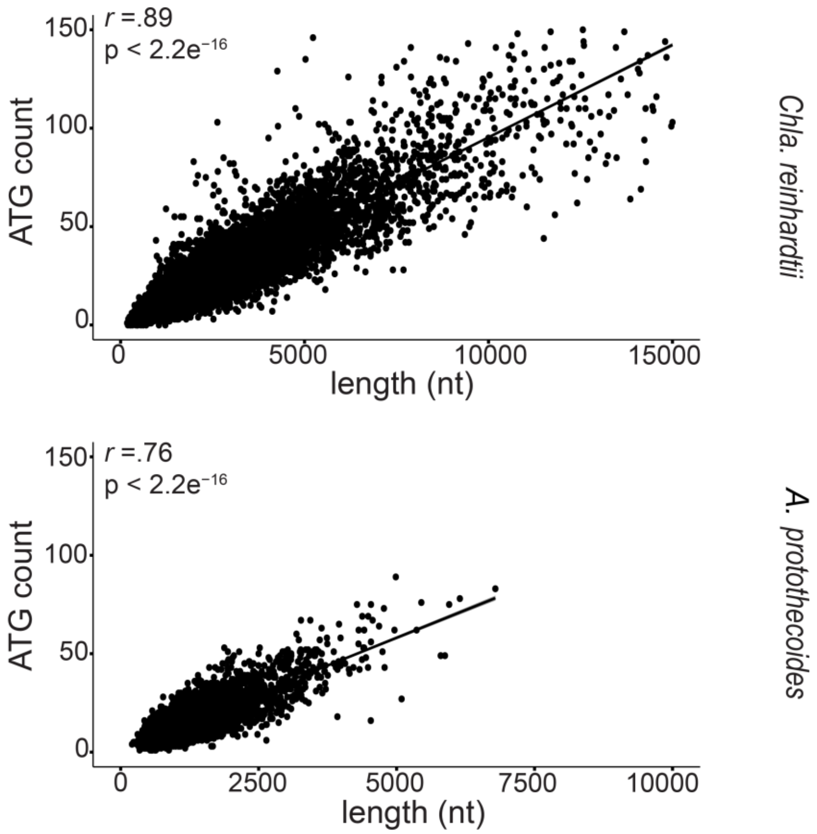


**Figure S2. Frequency of ATG codons correlates with increasing sequence length.**

The numbers of in and out of frame "ATG" sequences plotted against the sum of the lengths of the 5' UTR and the CDS for a set of monocistronic genes in the genomes of *A. protothecoides* (n = 7698) and *Chla. reinhardtii* (n =15,333). The line depicts a best fit linear regression. The correlation coefficient and *p*-value for significance are displayed at the top left of each graph.

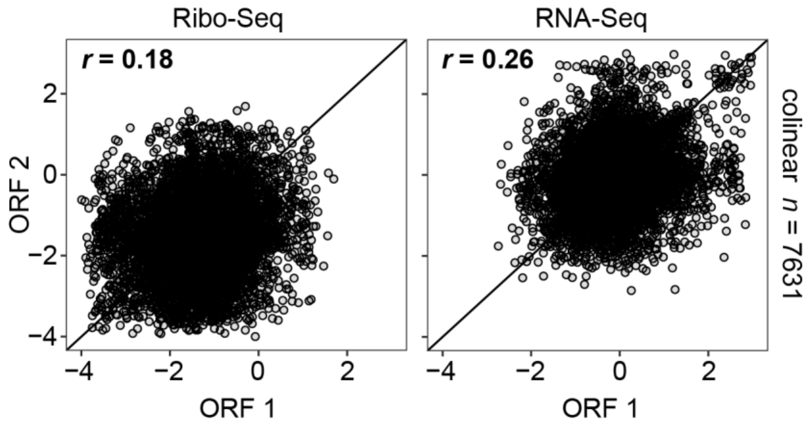


**Figure S3. Correlation of Ribo-Seq and RNA-Seq for the population of colinear genes**

Correlation analysis of the population of colinear genes ( *n* = 7631) in *Chla. reinhardtii*, which are defined here as adjacent genes on the same strand of the same chromosome with ≤10,000 nt between ORFs. A Pearsons correlation coefficient (*r*) was calculated as indicated. A diagonal line representing a perfect 1:1 correspondence is plotted for reference.

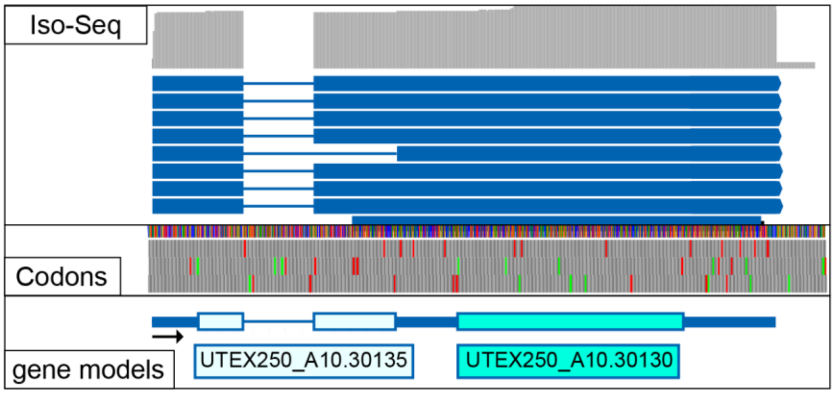


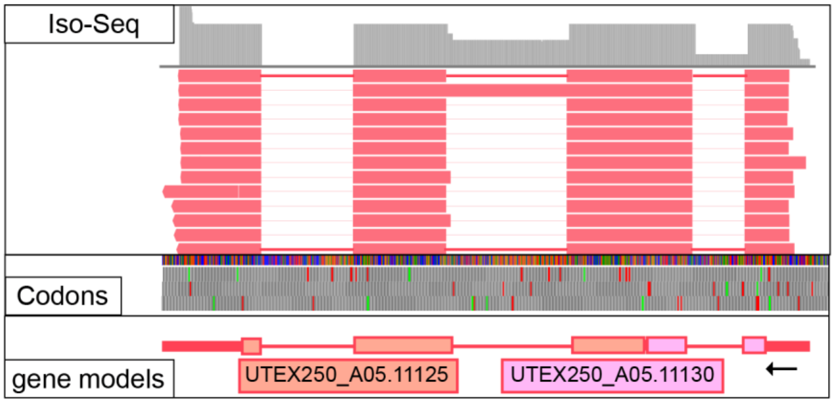


**Figure S4. Endogenous bicistronic loci for in vivo manipulation.**

IGV browser view of two endogenous bicistronic loci, predicted to encode ApTOM22_ApSDHAF (UTEX250_A10.30135/ UTEX250_A10.30130) and ApOST4_ApFAM32A (UTEX250_A05.11130/ UTEX250_A05.11130). For Iso-Seq, the forward (+) strand is shown in blue, and the reverse (-) strand is in pink. The black arrow indicates the 5' to 3' orientation of the strand. Coverage for each track is shown in gray. Stop and start codons are highlighted in red and green, all others are colored in grey. For gene models, a box indicates the exons of each ORF, a thick line indicates UTRs, and a thin line indicates introns.

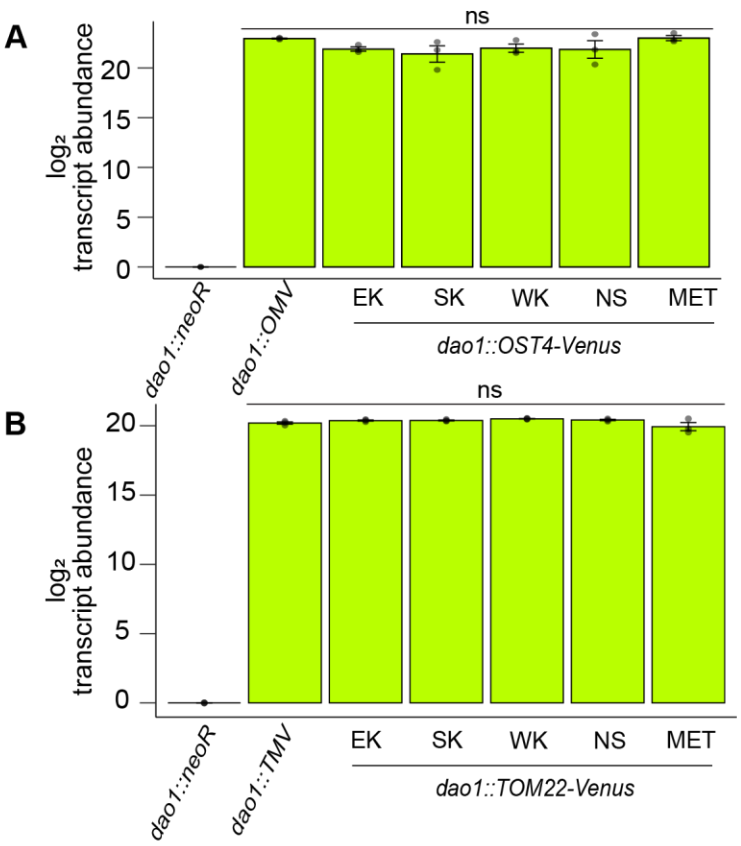


**Figure S5. Venus reporters show similar mRNA abundances**

Quantitative RT-qPCR analysis of **(A)** *OST4-Venus* and **(B)** *TOM22-Venus* reporter strains. Each datapoint represents the average of three biological replicates each averaged from technical duplicates. Error bars represent the standard deviation of the datapoints ($n$ = 3), with "ns" indicating a $p$-value greater than 0.05 from a one-way ANOVA statistical test.

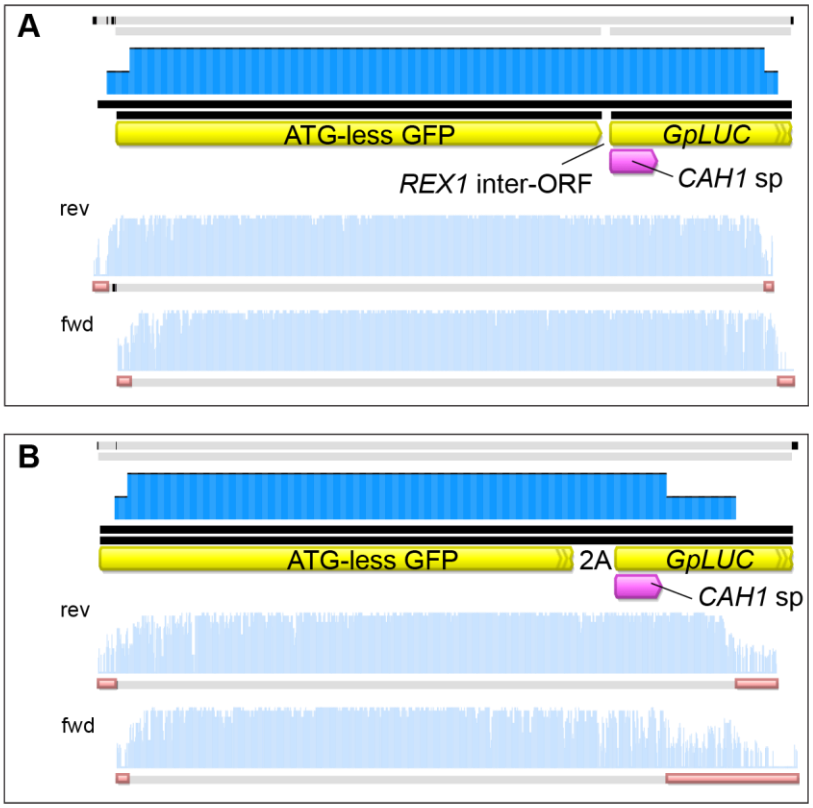


**Figure S6. Verification of synthetic bicistronic transcripts.**

Representative image of Sanger sequencing alignment of amplicons produced from RT-PCR reaction of GFP-LUC bicistronic strains. Yellow annotations denote the GFP and *Gaussia princeps* luciferase sequences. The pink annotation denotes the signal peptide from carbonic anhydrase 1 (*CAH1*). In **(A)** the region between the coding sequences refers to the 14bp inter-ORF from *ApREX1S-ApREX1B*. In **(B)** the region between the coding sequences refers to the 66bp 2A peptide sequence.

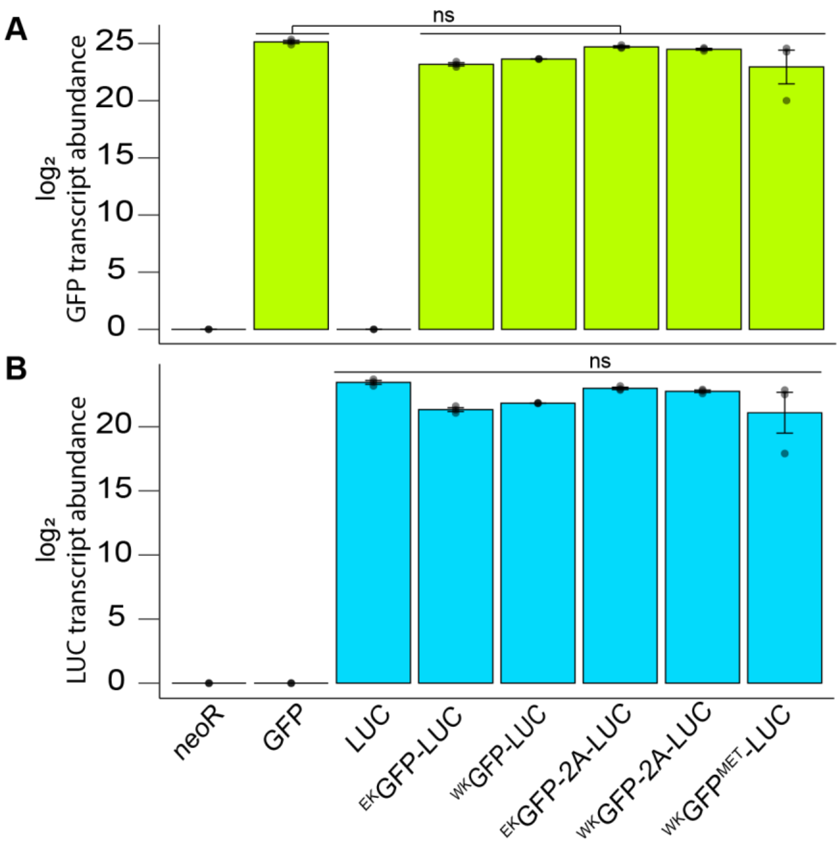


**Figure S7. The reporters are not differently expressed at the RNA level..**

Quantitative RT-qPCR analysis of the **(A)** GFP ORF and **(B)** Luciferase (LUC) ORF of GFP-LUC dual reporter strains. Each datapoint represents the average of three biological replicates measured in technical duplicates. Error bars represent the standard deviation of the datapoints (n = 3), with "ns" indicating a *p*-value greater than 0.05 from a one-way ANOVA statistical test.

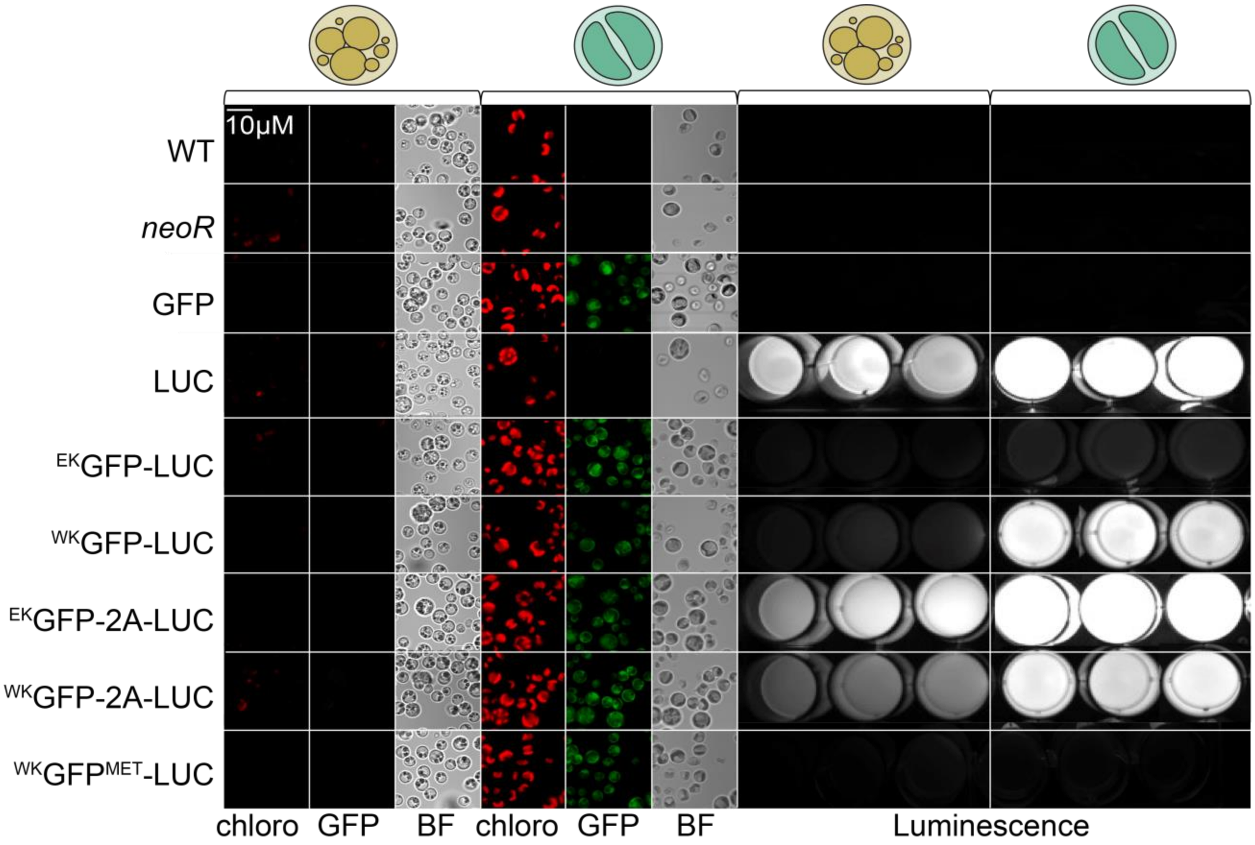


**Figure S8. Visualization of GFP and luciferase activity**

Representative images of confocal fluorescent microscopy and chemiluminescence images of dual reporter strains. For both heterotrophy (yellow cartoon) and photoautotrophy (green cartoon), panels display representative images of chlorophyll fluorescence (633nm excitation, 647-721nm emission), green fluorescent protein fluorescence (488nm excitation, 510-550nm emission), and the brightfield view respectively. The scale bar (white, top left) applies to all microscope images. Representative chemiluminescence of luciferase of three independent strains (*n* = 3) for cells grown in heterotrophy and photoautotrophy. At the time of imaging, luminescence was normalized to the $^{WK}$GFP-LUC strain.

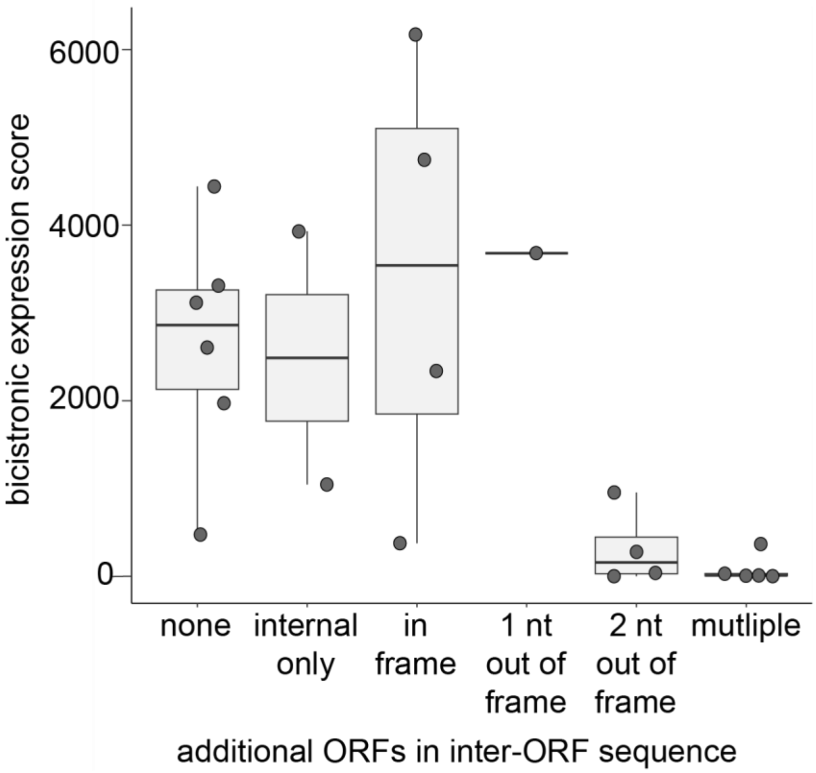


**Figure S9. The presence of other ORFs diminishes the bicistronic efficiency described in *Jacobebbinghaus et al*.**

Bicistronic expression scores of candidate sequences (identified as CS1 - CS22) from Jacobebbinghaus et al 2024. We evaluated each sequence for the presence of additional ORFs within the CS, and grouped them as follows: 1) those with no additional ORFs in the CS, 2) those with an ORF that begins and ends within the CS, 3) those with an ORF that is in-frame with ORF 2, 4) those with an ORF that is 1 nt out of frame relative to ORF 2, 5) those with an ORF that is 2 nt out of frame relative to ORF2, and 6) those with multiple and overlapping ORFs. Each of these is plotted here relative to the bicistronic expression score calculated to reflect the combined expression of both ORF 1 and ORF 2 (see Methods).

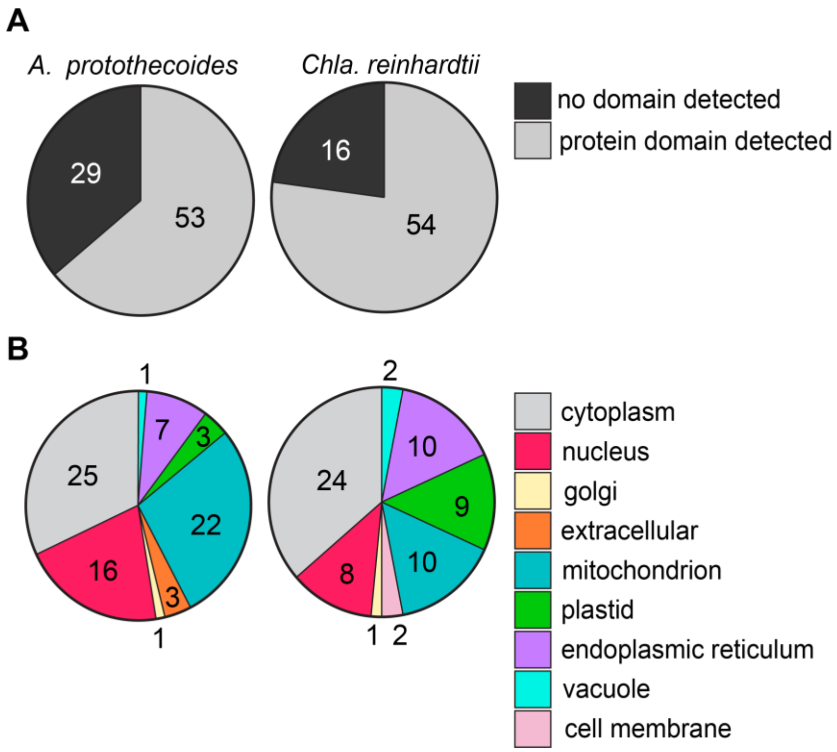


**Figure S10. Pfam and DeepLoc domain predictions of algal bicistronic loci.**

**(A)** Protein sequences from all bicistronic loci in this study were searched for conserved domains with InterPro scan. The number of proteins with an identified domain is presented as a pie chart for both *A. protothecoides* and *Chla. reinhardtii*. **(B)** Subcellular localization from all bicistronic loci in this study was predicted with DeepLoc 2.0. The top prediction of each protein is presented for both *A. protothecoides* and *Chla. reinhardtii*.
